# Multi-Omic and Spatial Profiling Defines Outcome-Associated Tumor States in Head and Neck Adenoid Cystic Carcinoma

**DOI:** 10.64898/2026.10.07.757413

**Authors:** Gopikrishnan Bijukumar, Kathryn J. Brayer, Anitha Vijaykumar, Elaine L. Bearer, David Y. Lee, Ryan K. Orosco, Scott A. Ness, Jeremy S. Edwards, Viswanathan Palanisamy

## Abstract

Head and neck adenoid cystic carcinoma (HNACC) shows highly variable clinical behavior that histopathological features only partly predict, and prognostic biomarkers for aggressive disease at diagnosis remain limited. Here, we show by integrating bulk, single-cell, and spatial transcriptomes from 25 HNACC tumors that poor survival is marked by profound loss of myoepithelial tumor cells and expansion of luminal tumor states. Both outcome groups share a myoepithelial-like origin, but aggressive tumors converge on an actively dividing, MYC/E2F-driven luminal state concentrated in discrete spatial niches. Aggressiveness therefore reflects where tumors end along a shared developmental path rather than a distinct lineage or altered immune infiltration. A four-gene classifier capturing this state predicted survival in an independent cohort, outperforming larger signatures and flagging aggressive subgroups in two others. These findings define a lineage proliferation axis linked to HNACC outcome, providing a basis for prospective risk stratification and proliferative-directed therapy.

## Introduction

Adenoid cystic carcinoma (ACC) is a rare malignancy arising predominantly from the secretory glands of the head and neck, accounting for approximately 1% to 2% of head and neck cancers and 10% of salivary gland malignancies (1–4). Although ACC often exhibits slow initial growth, it follows a paradoxically aggressive clinical course characterized by frequent perineural invasion, local recurrence, distant metastasis, and disease-specific mortality that may occur years after diagnosis (3,4). Current clinicopathologic features, including histologic grade, solid growth pattern, and stage, incompletely predict outcome, and patients with apparently similar tumors can experience markedly different clinical trajectories (5). Consequently, reliable biomarkers that can identify biologically aggressive disease at diagnosis remain a major unmet clinical need.

The molecular landscape of ACC is dominated by recurrent activation of the MYB transcriptional network (6). The hallmark genomic alteration is the MYB-NFIB fusion generated by t(6;9)(q22-23;p23-24) translocation, which drives aberrant transcriptional programs regulating proliferation, survival, and tumor progression (7–9). Tumors lacking MYB rearrangements frequently harbor analogous MYBL1 alterations, indicating that MYB and MYBL1 function as interchangeable oncogenic drivers within a common disease framework (6,10,11). Integrative genomic studies have further shown that these alterations converge on highly conserved transcriptional regulatory programs, establishing ACC as a transcription factor-driven malignancy with relatively low mutational burden. Notably, our earlier findings demonstrated that MYB and MYBL1 rearrangements define a shared oncogenic pathway in ACC, highlighting the central importance of transcriptional regulation in disease pathogenesis (6).

Beyond MYB-family activation, NOTCH signaling represents a second major pathogenic axis. Activating NOTCH1 alterations occur in approximately 8% to 26% of tumors and are enriched in recurrent and metastatic disease (9,12). Proteogenomic analyses subsequently identified two molecularly and clinically distinct ACC subtypes: ACC-I, characterized by MYC activation, NOTCH signaling, and poor prognosis, and ACC-II, characterized by TP63 expression and a more favorable clinical course (13). These studies have substantially advanced understanding of ACC biology, yet the cellular basis underlying these molecular programs and their relationship to patient outcome remain unclear.

Despite growing molecular insight, therapeutic progress has been limited. Surgical resection followed by radiotherapy remains the standard treatment for localized disease. Cytotoxic chemotherapy yields modest response rates, and targeted therapies directed against EGFR, cKIT, and VEGF have demonstrated limited clinical benefit (14,15). Although NOTCH inhibitors have shown activity in selected patients with NOTCH-activated recurrent or metastatic ACC, these agents remain investigational (16). Likewise, immune checkpoint inhibitors have generally underperformed, consistent with ACC’s immunologically cold phenotype (14,17). The lack of effective systemic therapies further underscores the need to identify biomarkers that distinguish aggressive disease and guide future therapeutic strategies.

Progress toward this goal has been complicated by the rarity of ACC and the scarcity of experimentally tractable models. Few authenticated ACC cell lines exist, genetically faithful animal models remain limited, and key MYB-dependent transcriptional circuits are often difficult to recapitulate outside patient tumors (18,19). As a result, patient-derived molecular datasets have become a critical resource for understanding ACC biology and discovering clinically relevant biomarkers. Reliable prognostic markers, however, remain elusive, and individual genomic alterations have not consistently stratified patients according to outcome (5).

Recent advances in transcriptomic technologies offer an opportunity to address this challenge. Bulk RNA sequencing can identify outcome-associated molecular programs but cannot distinguish whether such signals arise from differences in cellular composition, cell state, or tissue organization. Single-cell RNA sequencing resolves cellular heterogeneity and has revealed substantial variation in luminal and myoepithelial tumor-cell populations in ACC, including evidence that myoepithelial-derived NOTCH ligands may promote oncogenic signaling within luminal tumor cells (20,21). Spatial transcriptomics adds an additional layer of information by preserving the anatomical context of gene expression within intact tissue and enabling identification of region-specific transcriptional programs (22,23). Because each modality captures distinct aspects of tumor biology, integrating them provides an opportunity to define biologic features that would remain inaccessible through any single approach.

We hypothesized that clinical outcome in head and neck ACC (HNACC) is governed by coordinated changes in tumor-cell state, lineage composition, and spatial organization. To test this hypothesis, we integrated bulk RNA sequencing, single-cell RNA sequencing, and spatial transcriptomics from clinically annotated HNACC tumors stratified by long-term survival. This multimodal approach identified a reproducible outcome-associated lineage-proliferation axis characterized by loss of myoepithelial identity, expansion of proliferative luminal tumor states, activation of MYC/E2F- and G2/M-associated programs, and emergence of spatially organized proliferative niches. We further leveraged these findings to develop and externally validate a compact prognostic classifier, providing a biologically grounded framework for risk stratification and translational investigation in ACC.

## Results

### Cohort Design, Quality Control, and Integration of Single-Cell HNACC Datasets

To define the molecular determinants of clinical outcome in HNACC, we assembled a multi-modal transcriptomic cohort comprising 25 tumors arising from diverse anatomical sites (Fig. 1A). Patients were stratified according to overall survival into a good-outcome group (>5 years; n = 14, 56%) and a poor-outcome group (<2 years; n = 11, 44%). Three complementary transcriptomic platforms were analyzed, including bulk RNA sequencing (n = 20), single-cell RNA sequencing (n = 24), and spatial transcriptomics (n = 6), consisting of standard Visium (n = 4) and Visium HD (n = 2) datasets (Fig. 1B-C; Table 1).

**Fig. 1.**
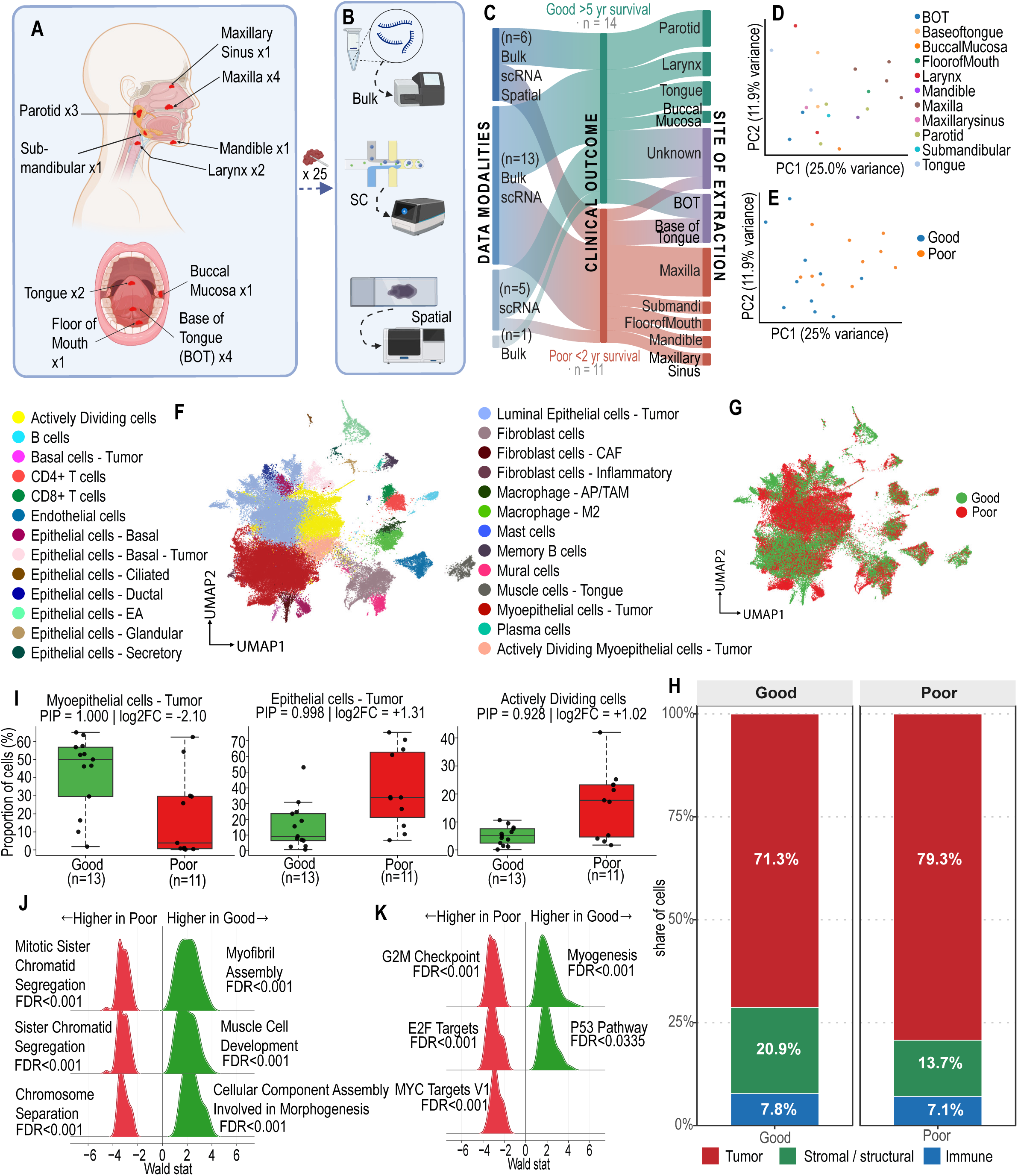
Multimodal cohort design and outcome-associated cellular shifts in HNACC. (A) Anatomical distribution of the 25 head and neck adenoid cystic carcinoma (HNACC) tumors included in the study. (B) Schematic overview of the multi-omic study design showing integration of bulk RNA sequencing, single-cell RNA sequencing (scRNA-seq), and spatial transcriptomic datasets. (C) Sankey plot summarizing sample allocation across transcriptomic platforms and clinical outcome groups defined by overall survival (<2 years, poor-outcome; >5 years, favorable-outcome). (D) Principal component analysis (PCA) of the bulk RNA-seq cohort colored according to anatomical site of origin. (E) The same PCA projection colored according to clinical outcome group. (F) Uniform Manifold Approximation and Projection (UMAP) of 95,441 single cells from 24 HNACC tumors following scVI-based batch correction and manual cell-type annotation. (G) The same integrated UMAP colored according to clinical outcome group. (H) Relative abundance of major cellular compartments after collapsing annotated cell populations into malignant, stromal/structural, and immune categories and aggregating samples by clinical outcome. (I) Cell populations exhibiting significant outcome-associated abundance differences identified by single-cell compositional analysis (scCODA). Mural cells served as the reference population. Box plots display sample-level cell-type proportions and significant posterior inclusion probability (credible at 5% estimated FDR (PIP ≥ 0.928) (J-K) Gene set enrichment analysis of pseudo bulked luminal epithelial tumor cells (LETCs) and actively dividing tumor cells (ADCs) comparing poor- and good-outcome tumors. Enrichment results are shown for (J) Gene Ontology Biological Process and (K) MSigDB Hallmark gene sets. Ridge plots depict the distribution of PyDESeq2 Wald statistics for genes in each pathway, based on 1,000 permutation-based enrichment analyses.

**Table 1.** Clinical characteristics of the 25 HNACC tumor samples and their multimodal allocations.

| Sample ID | Site of Extraction | Clinical Outcome | Clinical Categories | Bulk RNA Datasets | Single Cell RNA Datasets | Spatial Datasets |
| --- | --- | --- | --- | --- | --- | --- |
| A1197826 | Base of Tongue | < 2 years | Poor | Yes | Yes | No |
| A1386569 | Maxilla | < 2 years | Poor | Yes | Yes | Visium |
| A251463 | Submandibular Gland | < 2 years | Poor | Yes | Yes | No |
| A368571 | Maxilla | < 2 years | Poor | Yes | Yes | Visium HD |
| A45040 | Unknown | < 2 years | Poor | No | Yes | No |
| A644987 | Base of Tongue | < 2 years | Poor | Yes | Yes | No |
| A677996 | Maxilla | < 2 years | Poor | Yes | Yes | No |
| A7749534 | Floor of Mouth | < 2 years | Poor | Yes | Yes | Visium |
| A860040 | Mandible | < 2 years | Poor | Yes | Yes | No |
| A893263 | Maxillary Sinus | < 2 years | Poor | Yes | Yes | No |
| A90111 | Maxilla | < 2 years | Poor | Yes | Yes | No |
| A52398 | Unknown | > 3 years | Good | No | Yes | No |
| A166702 | Tongue | > 5 years | Good | Yes | Yes | Visium |
| A271554 | Parotid Gland | > 5 years | Good | Yes | Yes | No |
| A30281 | Unknown | > 5 years | Good | No | Yes | No |
| A317286 | Parotid Gland | > 5 years | Good | Yes | Yes | No |
| A50602 | Unknown | > 5 years | Good | No | Yes | No |
| A52190 | Unknown | > 5 years | Good | No | Yes | No |
| A672996 | Larynx | > 5 years | Good | Yes | Yes | No |
| A723677 | Base of Tongue | > 5 years | Good | Yes | Yes | Visium |
| A727826 | Larynx | > 5 years | Good | Yes | Yes | No |
| A742434 | Base of Tongue | > 5 years | Good | Yes | Yes | Visium<br>HD |
| A816076 | Tongue | > 5 years | Good | Yes | Yes | No |
| A863010 | Parotid Gland | > 5 years | Good | Yes | Yes | No |
| A917964 | Buccal Mucosa | > 5 years | Good | Yes | No | No |

We first examined the bulk RNA-seq cohort to determine whether global transcriptional variation was primarily associated with anatomical site or clinical outcome. Principal component analysis of the 3,000 most variable genes revealed no clear clustering according to anatomical site, indicating that HNACC maintains a largely conserved transcriptional architecture across distinct head and neck subsites (Fig. 1D). In contrast, tumors showed partial segregation according to outcome, suggesting the presence of survival-associated transcriptional programs despite substantial interpatient heterogeneity (Fig. 1E).

The single-cell cohort comprised 24 tumors, including 19 samples that overlapped with the bulk RNA-seq cohort. After sample-level quality control using Scanpy (24) consensus doublet removal and filtering, we retained 95,441 high-quality cells from an initial 103,487 captured cells. Initial dimensionality reduction demonstrated strong sample-specific clustering, consistent with batch-associated technical variation that obscured biological relationships across tumors (Fig. S1A). To generate a unified cellular atlas, datasets were integrated using single-cell variational inference (scVI) (25), which effectively reduced batch-associated variation while preserving biologically meaningful cellular structure (Fig. S1B). Unsupervised clustering followed by manual annotation identified the major malignant, epithelial, immune, stromal, and vascular cell populations present across HNACC tumors (Fig. 1F; Fig. S1C). Overlaying clinical outcome onto the integrated embedding showed broad representation of both survival groups across all major cellular compartments and suggested outcome-associated transcriptional differences within multiple cell populations (Fig. 1G). Collectively, these analyses established a high-quality single-cell atlas of HNACC spanning multiple anatomical sites and clinical outcomes. Notably, outcome-associated variation appeared to outweigh site-specific variation, providing a framework for investigating how cellular composition and tumor-cell states contribute to patient survival.

### Poor Clinical Outcome Is Associated with a Shift from Myoepithelial to Proliferative Luminal Tumor States

With an integrated single-cell atlas of HNACC established, we next investigated whether differences in cellular composition contribute to the disease’s survival-associated biology. For initial comparison, annotated cell populations were grouped into three broad compartments: tumor, stromal/structural, and immune cells. Poor-outcome tumors contained a greater proportion of malignant cells, whereas good-outcome tumors were enriched for stromal and structural populations. In contrast, the overall immune-cell fraction was similar between the two groups, suggesting that broad differences in immune infiltration are unlikely to explain the observed survival differences (Fig. 1H).

To identify specific cell populations associated with outcome, we performed compositional analysis across all 24 single-cell datasets using scCODA (26). Among the 26 annotated cell states, only three demonstrated significant outcome-associated abundance differences (FDR < 0.05) (Fig. 1I). The most pronounced changes occurred within the malignant compartment. Myoepithelial tumor cells were profoundly depleted in poor-outcome tumors (PIP = 1.000, log2FC = -2.10), declining from a median of 50.1% of malignant cells in good-outcome tumors to only 3.9% in poor-outcome tumors. Conversely, luminal epithelial tumor cells (LETCs) increased from 9.2% to 33.0% (PIP = 0.998, log2FC = 1.31), while actively dividing tumor cells (ADCs) expanded from 5.0% to 17.7% (PIP = 0.928, log2FC = 1.02). Across patients, myoepithelial and luminal tumor-cell proportions were strongly inversely correlated (Spearman ρ = -0.72, P = 1 × 10⁻⁴), indicating a coordinated shift in tumor-state composition rather than independent variation of individual cell populations.

We next asked whether expansion of the proliferative compartment reflected increased proliferation within luminal cells or expansion of the luminal lineage itself. The proportion of luminal cells classified as actively dividing was nearly identical between outcome groups (29.6% versus 29.3%; P = 0.772), indicating that the increased abundance of ADCs in poor-outcome tumors results primarily from expansion of the luminal compartment rather than a higher proliferation rate within individual luminal cells.

To define the transcriptional programs accompanying this lineage shift, we combined LETCs and ADCs into a unified luminal compartment and analyzed them using patient-level pseudobulk differential expression. This analysis identified 151 differentially expressed genes (FDR < 0.05, | log2FC| > 0.5), including 85 genes elevated in good-outcome tumors and 66 genes elevated in poor-outcome tumors. Pathway analysis revealed a highly directional pattern: nearly all significantly enriched pathways associated with poor-outcome were dominated by mitotic cell-cycle progression, chromosome segregation, and DNA replication programs (Fig. 1J-K), consistent with activation of a coordinated proliferative transcriptional state.

Because luminal cell abundance differed substantially between outcome groups, we performed multiple sensitivity analyses, including repeated cell down-sampling and count-thinning procedures. Gene-level effect estimates remained highly concordant across all analyses, and the same proliferation-associated pathways were consistently recovered (Fig. S1D), indicating that differences in cell abundance did not drive the observed transcriptional programs. Similar results were obtained when we repeated analyses across the entire single-cell cohort, independent of overlap with the bulk RNA-seq dataset. In these analyses, poor-outcome tumors remained enriched for proliferation- and cell-cycle-associated pathways. (Fig. S1E-G).

Taken together, these findings identify a previously unrecognized outcome-associated cellular transition in HNACC characterized by profound loss of the myoepithelial tumor compartment and expansion of a proliferative luminal tumor state. The consistency of this pattern across compositional and transcriptional analyses suggests that luminal lineage expansion is central to aggressive disease. We therefore next asked whether this shift in tumor-cell identity is accompanied by corresponding changes in intercellular communication within the tumor microenvironment.

### Intercellular Signaling Programs Distinguish Favorable and Unfavorable Clinical Outcomes in HNACC

The marked shift from myoepithelial to proliferative luminal tumor states prompted us to investigate whether these cellular changes were accompanied by remodeling of intercellular communication networks within the tumor microenvironment. Cell – cell communication analysis was implemented in Python using CellPhoneDB (27,28). Across the integrated single-cell dataset, malignant cells occupied the center of the inferred communication network and engaged extensively with both stromal and immune populations (Fig. 2A, E). Among all annotated cell populations, ADMEC (Actively Dividing Myoepithelial Cells)-tumor cells emerged as the dominant signaling hub, exhibiting extensive autocrine and paracrine interactions, particularly with cancer-associated fibroblasts (CAFs) and luminal epithelial tumor cells (LETCs) (Fig. S2A).

**Fig. 2.**
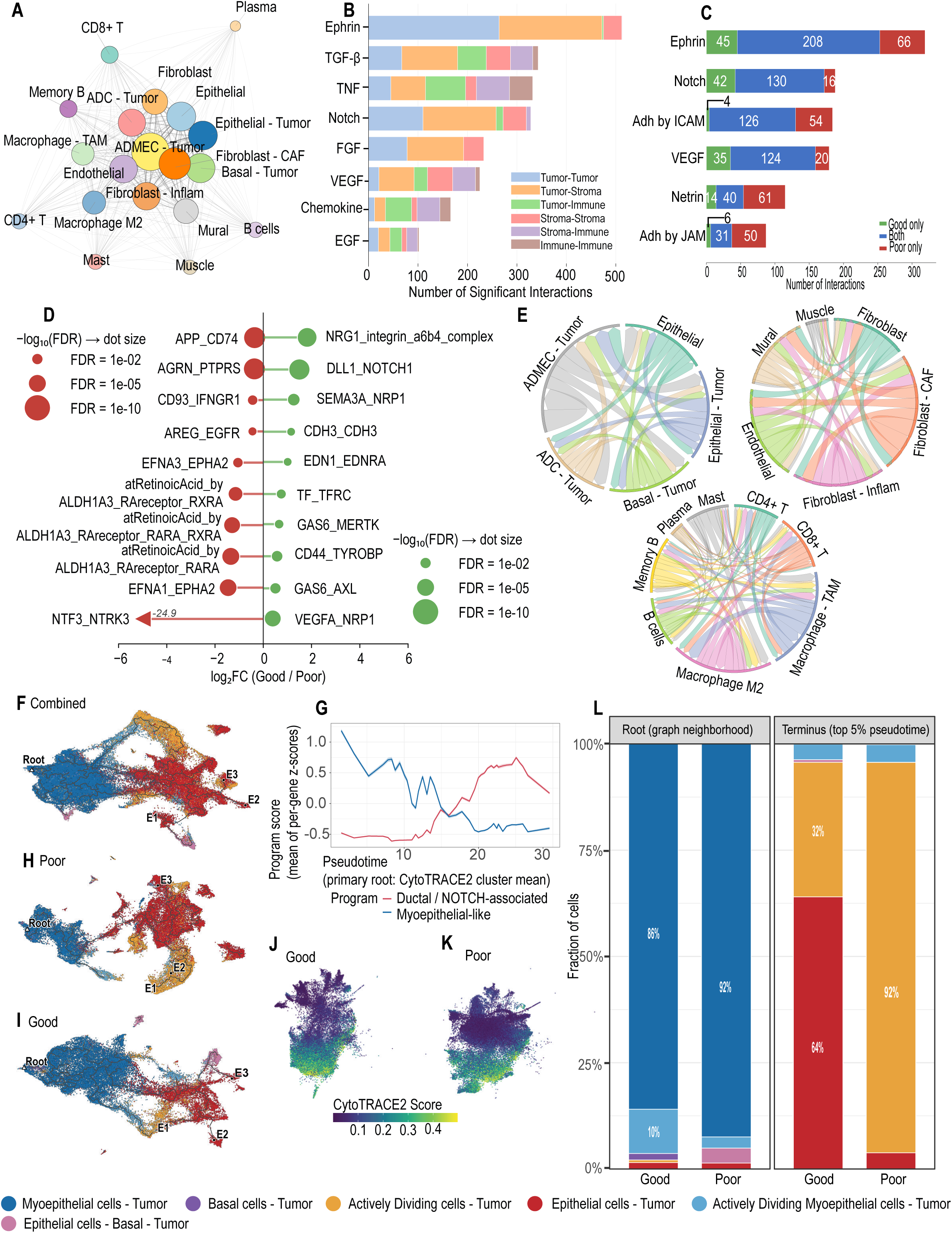
Outcome-associated cell-cell communication and developmental trajectories in HNACC. (A) Global CellPhoneDB-derived cell-cell communication network inferred across all 24 HNACC single-cell datasets. Node size is proportional to the number of significant ligand-receptor interactions involving each cell population. Edges represent significant communication events (P < 0.05). (B) Distribution of significant ligand-receptor interactions across major signaling pathways within the HNACC microenvironment. Ephrin, TGF-β, TNF, and NOTCH signaling represent dominant communication axes. (C) Number of significant ligand-receptor interactions detected exclusively in favorable-outcome tumors, exclusively in poor-outcome tumors, or shared between both outcome groups. (D) Differentially enriched ligand-receptor interactions between favorable- and poor-outcome cohorts following cell-number balancing across 15 cell populations. Positive values indicate enrichment in favorable-outcome tumors, whereas negative values indicate enrichment in poor-outcome tumors. The NTF3-NTRK3 interaction extends beyond the plotted x-axis range. (E) Chord diagrams summarizing significant interactions among tumor, stromal, and immune compartments. Ribbon width is proportional to the number of significant interactions between compartments. (F) Monocle3 pseudotime trajectory reconstructed from 71,984 malignant cells derived from 24 HNACC tumors. The trajectory was generated using highly variable genes and rooted independently of clinical outcome. (G) Myoepithelial-like and ductal/NOTCH-associated program activity across pseudotime. Program scores were calculated as the average standardized expression of genes within each signature. (H-I) Outcome-specific trajectory analyses. (H) Poor-outcome tumors. (I) Favorable-outcome tumors. Trajectories were reconstructed independently to evaluate differences in developmental endpoints while preserving outcome-blinded root selection. (J-K) CytoTRACE2-derived developmental potency scores calculated independently of trajectory inference for malignant cells from (J) favorable-outcome and (K) poor-outcome tumors. (L) Malignant cell-type composition at the trajectory root neighborhood and within the terminal 5% of pseudotime. Root states were enriched for myoepithelial tumor cells in both cohorts, whereas terminal states differed substantially, with poor-outcome tumors converging on a highly proliferative endpoint dominated by actively dividing tumor cells.

At the pathway level, Ephrin, TGF-β, TNF, and NOTCH signaling represented major communication axes within HNACC (Fig. 2B). Although most signaling pathways were detected across both clinical groups, their relative activity differed substantially according to outcome (Fig. 2C). Comparison of favorable- and poor-outcome tumors identified 123 of 227 evaluable ligand-receptor interactions with significantly different interaction strengths (FDR < 0.05), of which 96 were enriched in poor-outcome tumors (Fig. 2D). Poor-outcome tumors preferentially engaged signaling pathways associated with growth factor signaling, cellular migration, and neurotrophic activity, including AREG-EGFR, Ephrin-EPHA2, NTF3-NTRK3, AGRN-PTPRS, and NTN1-DSCAM interactions. The enrichment of neurotrophic signaling is particularly notable given the well-established neurotropism and propensity for perineural invasion that characterize clinically aggressive ACC.

In contrast, favorable-outcome tumors preferentially utilized signaling pathways associated with tissue organization, epithelial differentiation, and cellular homeostasis. Representative interactions included DLL1-NOTCH1, NRG1-ERBB3, NRG1-integrin α6β4, CDH3-CDH3, PDGFA-PDGFRA, and CXCL14-CXCR4 signaling. Importantly, most pathways remained detectable across both cohorts, suggesting that clinical outcome is associated primarily with differences in signaling intensity rather than complete gain or loss of specific communication programs.

Patient-level analyses further supported these observations. Eighty-three of the 123 differentially enriched interactions demonstrated directional concordance between pooled and patient-level analyses (Fig. S2B). Applying more stringent criteria based on patient-level effect direction, reproducibility across tumors, and stability after down-sampling yielded 46 high-confidence outcome-associated interactions, most of which remained enriched in poor-outcome tumors.

Among these, neurotrophic signaling emerged as one of the most reproducible features of aggressive disease. NTF3-NTRK3, NTF3-NTRK2, and NTN1-DSCAM interactions were consistently more frequent and exhibited greater interaction strengths in poor-outcome tumors (Fig. S2C-D). Although these interactions did not remain significant after multiple-testing correction at the patient level, their reproducible enrichment across complementary analytical approaches highlights neurotrophic signaling as a compelling hypothesis-generating feature of aggressive HNACC.

Collectively, these findings indicate that poor clinical outcome is associated not only with altered tumor-cell composition but also with substantially different rewiring of intercellular communication networks than seen in good-outcome tumors. Poor-outcome tumors preferentially engage growth factor, migratory, and neurotrophic signaling programs, whereas favorable-outcome tumors retain communication networks associated with cellular differentiation and tissue organization. These observations raised the possibility that the two clinical groups occupy distinct developmental states, prompting us to examine malignant-cell differentiation trajectories.

### Trajectory Analysis Reveals Divergence Toward a Proliferative Luminal Terminal State in Poor-Outcome HNACC

The outcome-associated differences in cellular composition and intercellular signaling suggested that favorable- and poor-outcome tumors may represent distinct points along a common developmental continuum. To investigate this possibility, we reconstructed malignant-cell trajectories using 71,984 tumor cells derived from 24 HNACC specimens using CytoTrace2 (29) and Monocle3 (30) in R.

Trajectory analysis identified a continuous developmental landscape connecting the major malignant populations, with myoepithelial tumor cells occupying the root of the trajectory and luminal populations residing along progressively differentiated states (Fig. 2F). Independent assessment of developmental potential using CytoTRACE2 supported this architecture, identifying myoepithelial and basal tumor cells as the most developmentally potent malignant populations (Fig. 2J-K). As pseudotime progressed, myoepithelial-associated programs progressively declined, whereas ductal and NOTCH-associated programs increased, consistent with differentiation from a myoepithelial-like state toward a luminal tumor identity (Fig. 2G; Fig. S3F).

The inferred developmental hierarchy proved highly robust. Multiple outcome-blinded rooting strategies consistently identified myoepithelial tumor cells as the trajectory origin (Fig. 2F; Fig. S3A). Similarly, analyses performed independently within favorable- and poor-outcome tumors recovered the same overall architecture, with the root neighborhood remaining strongly enriched for myoepithelial tumor cells in both cohorts (85.8% and 92.5%, respectively; Fig. 2H-I; Fig S3B-C).

Despite sharing a common developmental origin, the two outcome groups diverged strikingly at the terminal stages of differentiation. Examination of cells occupying the terminal 5% of pseudotime revealed that favorable-outcome tumors retained a mixed luminal endpoint composed predominantly of luminal epithelial tumor cells (64%) together with actively dividing tumor cells (32%). In contrast, poor-outcome tumors converged almost exclusively on a highly proliferative endpoint dominated by actively dividing tumor cells (92%), with only a small residual population of differentiated luminal epithelial tumor cells (4%) (Fig. 2L).

To further characterize these divergent terminal states, we examined pathway activity along the developmental trajectory. NOTCH signaling increased progressively along pseudotime and reached maximal activity during acquisition of luminal identity (Fig. S3F). Importantly, malignant cells from poor-outcome tumors consistently exhibited greater NOTCH pathway activity than corresponding cell populations from favorable-outcome tumors (Fig. S3G-H), suggesting enhanced activation of luminal differentiation and proliferation-associated programs in aggressive disease. In contrast, EMT-associated programs were detected in both clinical groups but occurred in distinct cellular contexts and did not consistently track with outcome (Fig. S3D-H).

Together, these findings support a developmental model in which HNACC tumors originate from a shared myoepithelial-like state but diverge toward distinct terminal phenotypes. Favorable-outcome tumors retain a more differentiated luminal endpoint, whereas poor-outcome tumors preferentially accumulate within a highly proliferative luminal terminal state characterized by enhanced NOTCH activity. When considered alongside the compositional, transcriptional, and signaling analyses, these observations suggest that aggressive HNACC is driven not simply by loss of myoepithelial identity, but by progression toward a proliferative luminal state that may represent a clinically actionable therapeutic vulnerability.

### Outcome-Associated Transcriptional Programs Reveal Molecular Features of Aggressive ACC

Having identified outcome-associated differences in tumor-cell composition, intercellular communication, and developmental state at single-cell resolution, we next asked whether these biological distinctions were reflected at the bulk transcriptomic level. We performed differential expression analysis on 20 ACC tumors comprising 10 good-outcome and 10 poor-outcome cases using PyDESeq2 (31). A total of 1,191 genes were differentially expressed between outcome groups (FDR < 0.05, |log2FC| ≥ 0.5), including 516 genes enriched in good-outcome tumors and 675 genes enriched in poor-outcome tumors.

Among the most significantly altered transcripts, DSCAM was strongly upregulated in poor-outcome tumors, whereas PCAT14 was among the most highly expressed genes in good-outcome disease (Fig. 3A, S4A). Several genes previously implicated in ACC biology also demonstrated outcome-associated expression patterns. Good-outcome tumors showed higher expression of MUC12 and KDM6A, whereas poor-outcome tumors exhibited increased expression of oncogenic regulators including MYC, FGFR3, and TP53 (Fig. 3A). Examination of a curated set of 103 published ACC-associated genes further supported selective transcriptional remodeling between survival groups. Of the 103 ACC associated genes, 100 were represented in our dataset (Supplementary Table 2) out of which 31 were significantly differentially expressed, whereas the majority remained unchanged (Fig. 3B), indicating that poor clinical outcome is associated with activation of specific biological programs rather than broad transcriptomic divergence.

**Fig. 3.**
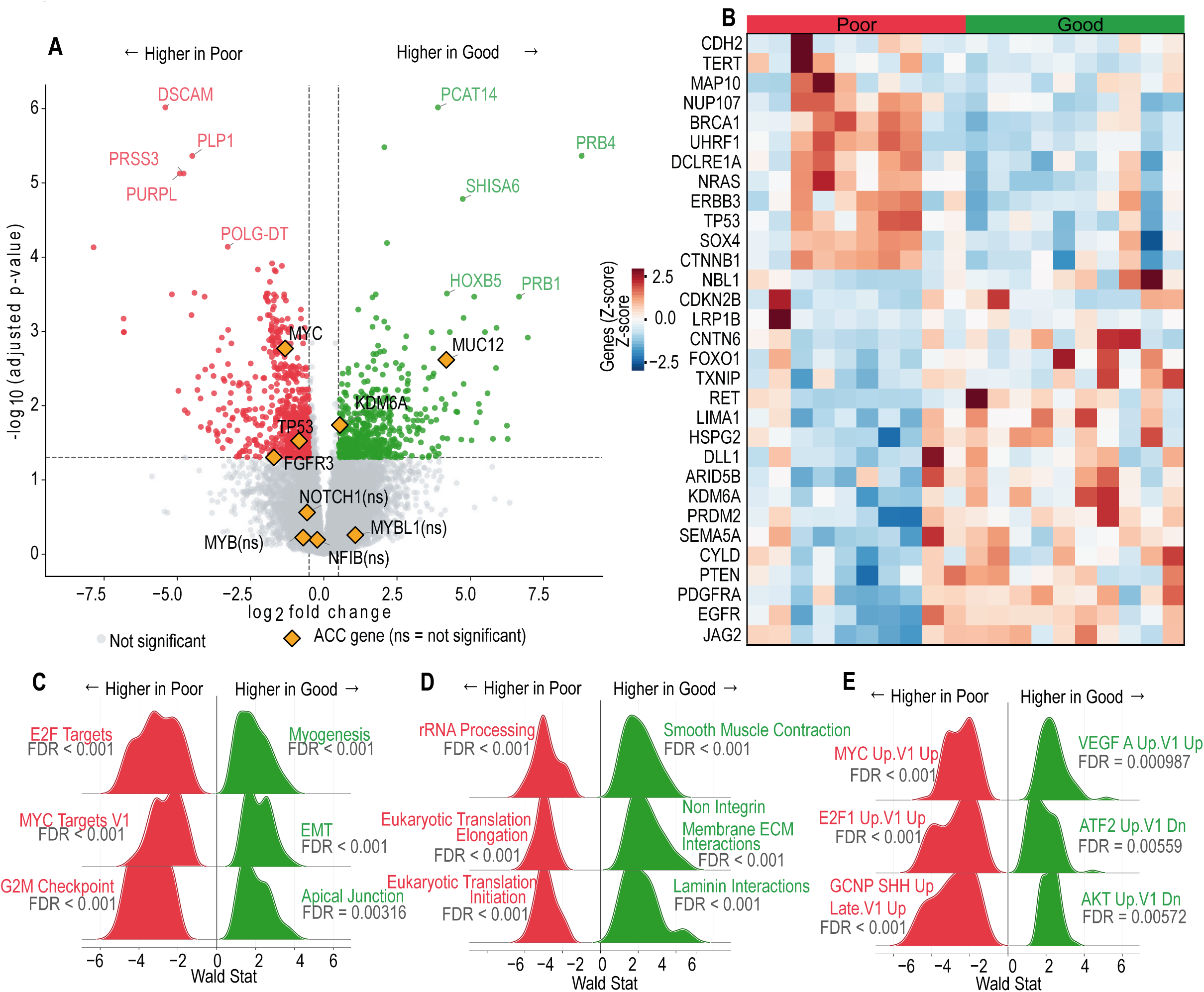
Bulk transcriptomic profiling identifies a MYC/E2F-driven proliferative program associated with poor clinical outcome in HNACC. (A) Differential expression analysis of the bulk RNA-seq cohort (n = 20; 10 good-outcome and 10 poor-outcome tumors) performed using PyDESeq2. Vertical and horizontal dashed lines indicate significance thresholds (FDR < 0.05 and |log2FC| ≥ 0.5). Positive log2 fold changes indicate genes enriched in favorable-outcome tumors, whereas negative log2 fold changes indicate genes enriched in poor-outcome tumors. Selected ACC-associated genes are highlighted. Genes not meeting significance thresholds are denoted as *ns*. (B) Heatmap showing row-scaled expression of 31 published ACC-associated genes that were significantly differentially expressed (FDR < 0.05) between outcome groups. Samples are arranged according to clinical outcome. (C) Gene set enrichment analysis (GSEA) of the bulk RNA-seq cohort against the MSigDB Hallmark collection using genes ranked by the PyDESeq2 Wald statistic. Enrichment reveals activation of MYC, E2F, G2/M checkpoint, and proliferation-associated programs in poor-outcome tumors, whereas good-outcome tumors are enriched for extracellular matrix and differentiation-associated programs. (D) GSEA was performed against the Reactome pathway collection. Poor-outcome tumors exhibit enrichment of pathways involved in translation, ribosome biogenesis, RNA processing, and cell-cycle progression, consistent with increased biosynthetic and proliferative activity. All enrichment analyses were performed using preranked GSEA with 1,000 permutations. Normalized enrichment scores are shown for pathways meeting statistical significance (FDR < 0.05). (E) GSEA was performed against the MSigDB Oncogenic Signatures collection. Poor-outcome tumors demonstrate enrichment of MYC- and E2F-associated transcriptional programs.

Pathway-level analyses using GSEAPy (32) revealed a remarkably consistent pattern across all enrichment frameworks. A total of 1,617 gene sets were significantly enriched (FDR < 0.05), with the overwhelming majority associated with poor clinical outcome (Fig. 3C-E). Across multiple pathway collections (MSigDB, Reactome, Gene Ontology, and KEGG), poor-outcome tumors consistently exhibited evidence of heightened proliferative activity. Hallmark analyses demonstrated strong enrichment of E2F targets, MYC targets, and G2/M checkpoint programs, whereas oncogenic-signature analyses independently identified activation of MYC- and E2F1-associated transcriptional networks (Fig. 3C, E).

This proliferative phenotype extended beyond cell-cycle regulation alone. Reactome analyses showed enrichment of translation initiation and elongation machinery, ribosome biogenesis, and RNA-processing pathways (Fig. 3D). Concordant results from Gene Ontology and KEGG collections revealed enrichment of DNA replication, chromosome segregation, replication licensing, and mRNA export programs (Fig. S4B-E). Together, these findings indicate that poor-outcome tumors could be characterized by increased biosynthetic capacity coupled with active cell-cycle progression.

In contrast, good-outcome tumors were enriched for pathways associated with extracellular matrix organization, cell adhesion, myogenesis, and tissue differentiation (Fig. 3C-E; Fig. S4B-E), consistent with preservation of a more differentiated and structurally organized tumor phenotype. Notably, these pathway-level patterns closely mirrored those identified in the single-cell analyses, where good-outcome tumors were enriched for myoepithelial and stromal programs, whereas poor-outcome tumors were dominated by proliferative luminal states.

Taken together, these data establish MYC/E2F-driven proliferation as a defining molecular feature of aggressive ACC. More importantly, they provide independent bulk transcriptomic evidence that poor clinical outcome is associated with activation of coordinated cell-cycle, translational, and biosynthetic programs that parallel the expansion of proliferative luminal tumor states observed at single-cell resolution. The remarkable agreement between bulk and single-cell analyses suggests that this proliferative luminal program represents a fundamental feature of aggressive HNACC biology.

### Co-expression Network Analysis Independently Recapitulates Outcome-Associated Proliferative and Stromal Programs

To determine whether the transcriptional differences identified by differential expression analysis reflected coordinated biological networks rather than isolated gene-expression changes, we performed weighted gene co-expression network analysis (WGCNA) (33) on the bulk RNA-seq cohort in R. Following quality-control filtering and removal of one outlier sample (A816076), network construction was performed at a soft threshold value of 14, using 19 tumors and 16,639 expressed genes. A signed co-expression network identified 23 gene modules, three of which demonstrated significant associations with clinical outcome (Fig. 4A; Fig. S5A–C).

**Fig. 4.**
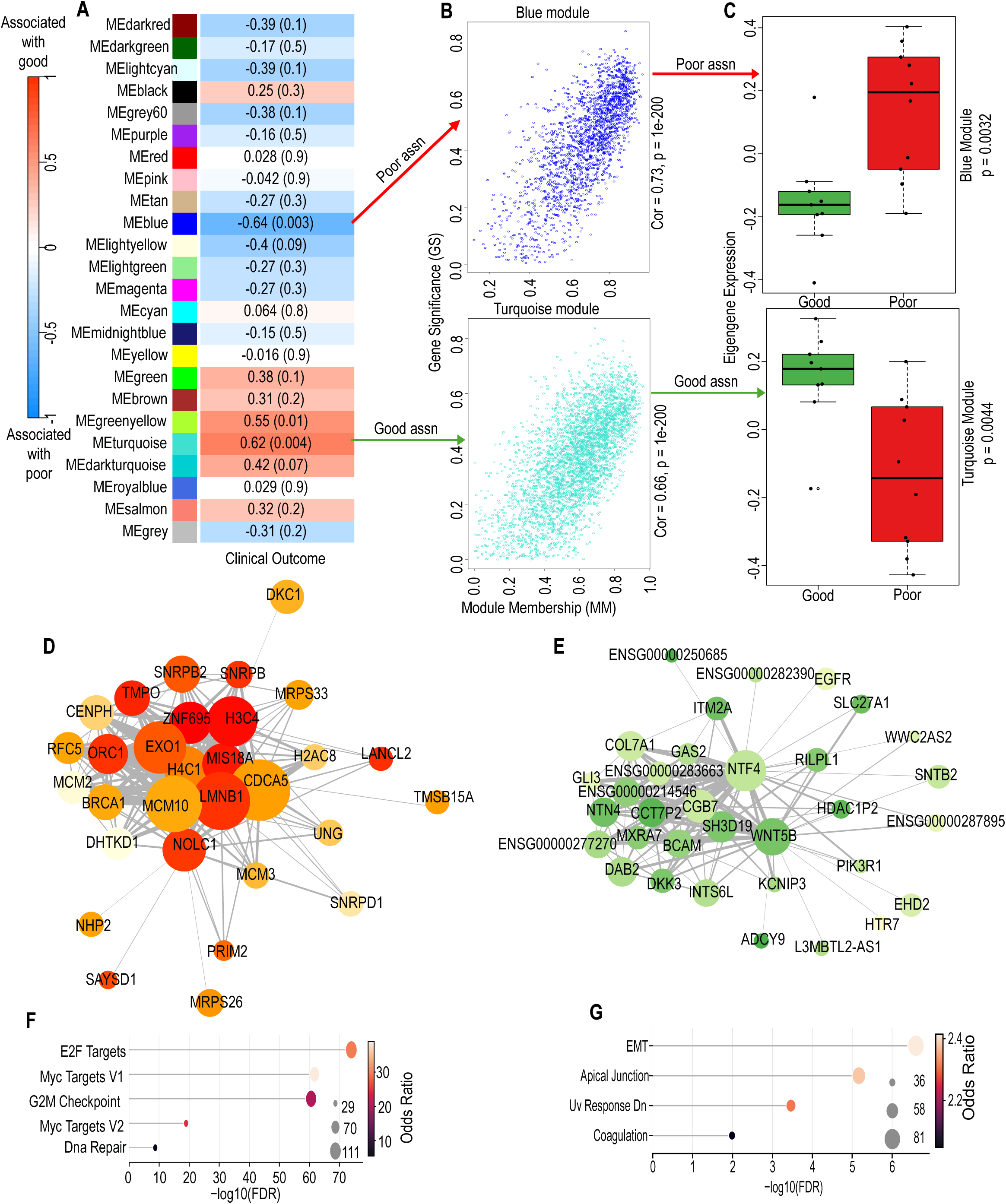
Coexpression network analysis independently identifies proliferative and stromal programs associated with clinical outcome in HNACC. (A) Module-trait relationships derived from weighted gene co-expression network analysis (WGCNA) performed on 19 bulk RNA-seq tumors and 16,639 expressed genes following removal of one outlier sample. Cells display Pearson correlation coefficients between module eigengenes and clinical outcome (favorable-outcome = 1, poor-outcome = 0), with corresponding asymptotic P values shown in parentheses. Modules significantly associated with outcome are highlighted. (B) Relationships between gene significance (GS) and module membership (MM) for the outcome-associated blue and turquoise modules. Genes exhibiting high GS and MM values were considered candidate hub genes. (C) Distribution of module eigengene expression in outcome-associated modules stratified by clinical outcome group. (D-E) Network representation of hub-gene architecture within the poor-outcome blue module (D) and good-outcome turquoise module (E). Networks were constructed from topological overlap matrices (TOMs), retaining highly connected edges. Node size and color are proportional to intramodular connectivity (kWithin), highlighting genes occupying central positions within each network. (F-G) Over-representation analysis (ORA) of genes belonging to the poor-outcome blue module (F) and good-outcome turquoise module (G) using the MSigDB Hallmark collection. The poor-outcome module is enriched for MYC targets, E2F signaling, G2/M checkpoint regulation, DNA replication, and proliferation-associated pathways, whereas the good-outcome module is enriched for extracellular matrix organization and epithelial to mesenchymal transition programs.

Among these, the blue module showed the strongest negative correlation with outcome and was therefore associated with poor prognosis (r = -0.64, P = 0.003), whereas the turquoise module demonstrated a strong positive association with favorable outcome (r = 0.62, P = 0.004) (Fig. 4A). A third module (greenyellow) was also associated with favorable outcome (r = 0.55, P = 0.01) but exhibited weaker biological coherence and was not pursued further. Consistent with these associations, eigengene expression for both the blue and turquoise modules differed significantly between outcome groups (Fig. 4C).

To identify genes most strongly associated with these outcome-linked networks, we examined the relationship between module membership and gene significance (Fig. 4B). The poor-outcome blue module was centered on hub genes including NOLC1, EXO1, and CDCA5, whereas the good-outcome turquoise module was driven by genes including NTF4, WNT5B, and BCAM. Network visualization revealed dense intramodular connectivity surrounding these hub genes, suggesting that they occupy central regulatory positions within their respective transcriptional programs (Fig. 4D-E).

We next investigated the biological processes represented within these outcome-associated networks. Consistent with the bulk differential expression analyses, the poor-outcome blue module was strongly enriched for MYC targets, E2F signaling, G2/M checkpoint regulation, DNA replication, and DNA repair pathways (Fig. 4F). These findings indicate activation of a coordinated proliferative and biosynthetic program in poor-outcome tumors. In contrast, the good-outcome turquoise module showed enrichment for extracellular matrix organization, collagen-associated pathways, coagulation, apical junction, and stromal differentiation programs, consistent with preservation of a more structured and differentiated tumor microenvironment (Fig 4G; Fig. S5D).

Importantly, these network-derived programs closely mirrored the patterns identified through both bulk transcriptomic and single-cell analyses. Across all three analytical approaches, poor-outcome tumors were consistently characterized by activation of MYC/E2F-driven proliferative programs, whereas good-outcome tumors retained stromal, extracellular matrix, and differentiation-associated signatures. The convergence of these independent analyses strongly suggests that expansion of a proliferative luminal state and loss of myoepithelial-associated programs represent fundamental features of aggressive HNACC biology rather than dataset-specific observations.

### Spatial Transcriptomic Profiling Validates Outcome-Associated Differences in Tissue Architecture

Having identified outcome-associated differences in cellular composition, developmental state, and transcriptional programs, we next asked whether these features were preserved within intact tumor tissue. To address this, we performed spatial transcriptomic profiling of six HNACC specimens using the 10x Genomics Visium platform using Giotto in R (34), including four standard-resolution samples (two favorable-outcome and two poor-outcome) and two Visium HD samples (one favorable-outcome and one poor-outcome).

Cell-type deconvolution was performed using Dampened Weighted Least Squares (DWLS) (35,36). Our single cell atlas developed in this study was used as reference for deconvolution. This revealed marked differences in tissue composition between outcome groups (Fig. 5A). Poor-outcome tumors were dominated by malignant populations, with tumor cells accounting for 49.1% and 57.9% of deconvoluted profiles, compared with only 15.0% and 9.5% in good-outcome tumors. This distinction was driven primarily by expansion of actively dividing tumor cells (ADCs), which comprised 38.9% and 36.1% of cells in poor-outcome tissues but only 5.7% and 0.03% in good-outcome samples. In contrast, mural and muscle-like populations were markedly enriched in good-outcome tumors, representing 9.9% and 44.8% of cells compared with only 0.9% and 1.4% in poor-outcome tissues. Normal epithelial populations were similarly more abundant in good-outcome specimens, whereas fibroblasts remained a major component of all tumors and showed relatively stable abundance across outcome groups. Importantly, these spatially inferred cellular compositions was in accordance with the compositions observed in the single-cell cohort (Fig. S6A).

**Fig. 5.**
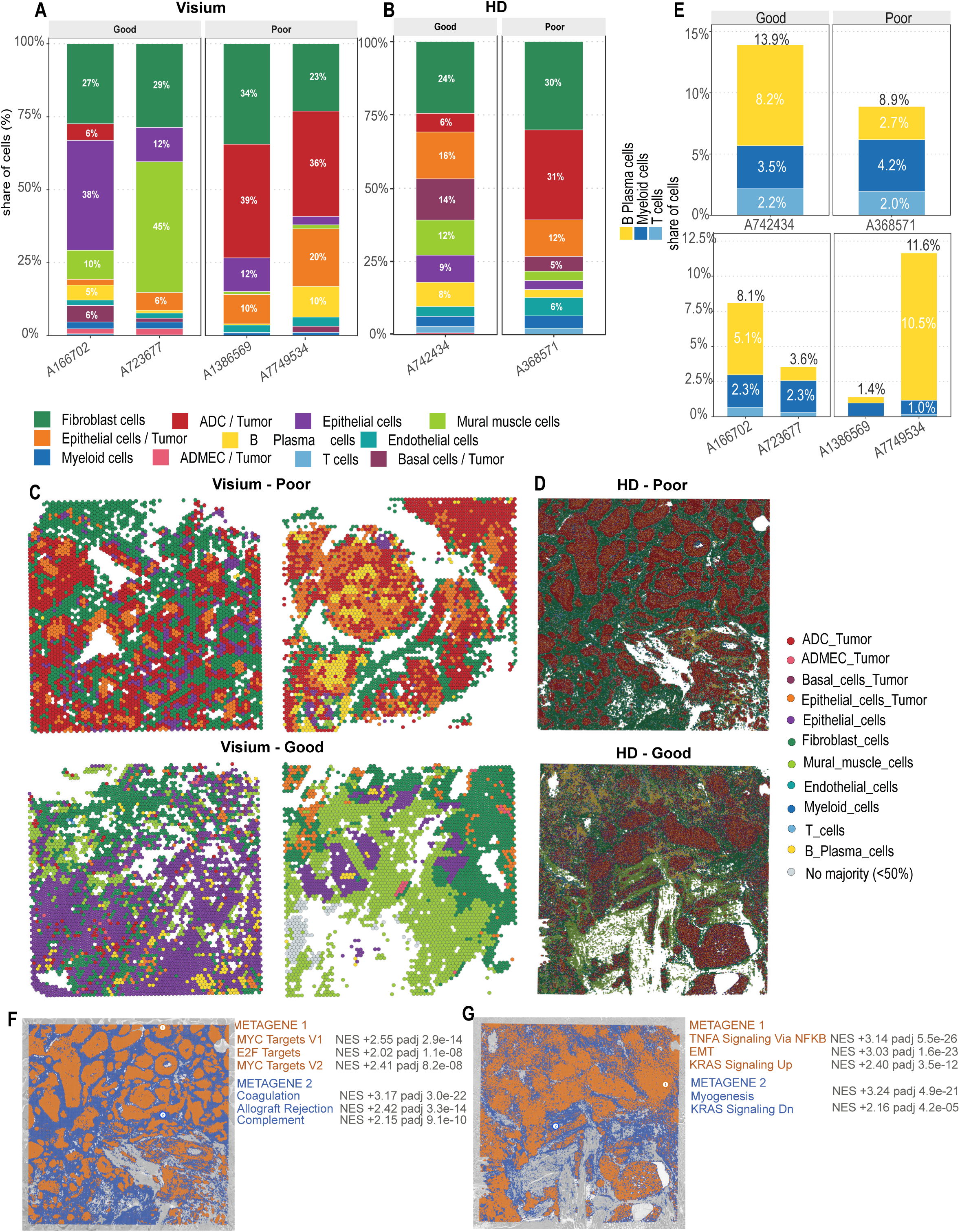
Spatially resolved cellular composition and transcriptional organization of favorable- and poor-outcome HNACC. (A) SpatialDWLS-based cell-type deconvolution of four standard-resolution Visium tissue sections (55-µm spot resolution) using the 24-sample single-cell RNA-seq atlas as the reference. Poor-outcome tumors are enriched for malignant and actively dividing tumor-cell populations, whereas favorable-outcome tumors are enriched for mural, muscle-like, and epithelial populations. (B) SpatialDWLS deconvolution of Visium HD samples generated at 2-µm resolution and aggregated into 16-µm bins. Higher spatial resolution reveals additional cellular populations, including basal tumor cells, while confirming enrichment of actively dividing tumor cells in poor-outcome tumors. (C-D) Spatial distribution of dominant cell populations following deconvolution. Each spatial location is assigned according to the cell type comprising more than 50% of the inferred composition. (C) Standard Visium sections and (D) Visium HD sections. Poor-outcome tumors display compact tumor-rich nests embedded within fibroblast-rich stroma, whereas favorable-outcome tumors exhibit a more heterogeneous and intermixed tissue architecture. (E) Relative abundance of immune-cell populations inferred by spatial deconvolution. Immune-cell composition varies between individual specimens but does not demonstrate a consistent association with clinical outcome. (F-G) Spatial metagene territories identified in Visium HD tumors using the 250 most spatially variable genes. Metagene modules were generated by Ward-linkage hierarchical clustering and projected onto tissue sections. (F) Poor-outcome tumor showing a dominant proliferative territory enriched for MYC- and E2F-associated programs. (G) Favorable-outcome tumor showing spatially distinct transcriptional territories enriched for contractile, stromal, and differentiation-associated programs. Representative MSigDB Hallmark pathways significantly enriched within each metagene territory are indicated.

The higher spatial resolution afforded by Visium HD further reinforced these findings. Basal tumor cells, which were only minimally resolved at standard Visium resolution, accounted for 14.0% of cells in the good-outcome sample but only 5.1% in the poor-outcome sample (Fig. 5B). Consistent with the standard-resolution analyses, fibroblast abundance remained relatively stable, whereas ADCs remained strongly enriched in poor-outcome tissues.

Beyond cellular composition, the spatial organization of the tumor microenvironment differed substantially between outcome groups. Standard-resolution Visium data revealed a nested tumor architecture in poor-outcome tumors, whereas good-outcome tumors exhibited a more heterogeneous, intermixed cellular landscape (Fig. 5C). These differences were even more apparent in the Visium HD datasets. Poor-outcome tumors were characterized by compact tumor nests embedded within fibroblast-rich stromal regions, whereas good-outcome tumors displayed a more complex architecture composed of intermixed epithelial, mural, muscle-like, fibroblast, and immune-cell territories (Fig. 5D).

In contrast to the pronounced differences observed within the malignant and stromal compartments, immune-cell distributions showed no consistent association with clinical outcome. Total immune-cell abundance varied across specimens and spatial platforms, with B/plasma-cell populations accounting for most of the observed variation (Fig. 5E). These findings are consistent with the single-cell analyses, which likewise suggested that broad differences in immune-cell abundance are not a primary determinant of outcome in HNACC.

Taken together, spatial transcriptomic profiling independently showed the cellular states identified through single-cell sequencing and demonstrated that poor-outcome HNACC is characterized by expansion of proliferative malignant populations organized into discrete tumor nests. In contrast, good-outcome tumors retain a more heterogeneous tissue architecture enriched for mural, epithelial, and stromal populations. These findings indicate that the biologic programs associated with clinical outcome are reflected not only in cellular composition but also in the spatial organization of the tumor microenvironment.

### Spatially Resolved Metagene Mapping Reveals Proliferative Niches in Aggressive ACC

Having established that favorable- and poor-outcome tumors exhibit distinct cellular compositions and tissue architectures, we next asked whether the transcriptional programs associated with clinical outcome were similarly organized into discrete spatial territories within intact tumor tissue. Spatial metagene analysis identified transcriptionally coordinated domains across all Visium samples, revealing substantial intratumoral organization. Although no individual pathway was exclusive to either outcome group, the composition and relative abundance of spatial programs differed markedly by clinical outcome.

Poor-outcome tumors consistently contained territories enriched for proliferative and tumor-promoting programs, including MYC targets, E2F targets, EMT -associated signaling, complement activation, coagulation pathways, and TNFα/NF-κB signaling (Fig. S6B). These spatial domains closely recapitulated the proliferative programs identified independently through single-cell transcriptomics, bulk RNA sequencing, and co-expression network analyses, providing spatial evidence that the aggressive transcriptional state observed across platforms is concentrated within localized tumor regions rather than uniformly distributed throughout the tissue.

In contrast, good-outcome tumors displayed considerably greater transcriptional heterogeneity. Distinct spatial territories were enriched for oxidative phosphorylation, respiratory electron transport, protein secretion, keratinization, myogenesis, and extracellular matrix organization programs (Fig. S6C). Although EMT-associated signatures were also detected in good-outcome tumors, they occurred within regions enriched for differentiated stromal and contractile programs rather than within highly proliferative tumor territories. These findings are consistent with preservation of a more differentiated and structurally organized tumor microenvironment.

The higher-resolution Visium HD datasets further reinforced these observations. In the poor-outcome tumor, the dominant metagene territory occupied approximately half of all analyzed bins and was strongly enriched for MYC- and E2F-associated transcriptional programs, recapitulating the proliferative state identified at standard Visium resolution (Fig. 5F). In contrast, the good-outcome tumor demonstrated clear spatial segregation of biologically distinct programs. One territory was enriched for myogenesis and contractile signatures, consistent with the increased abundance of mural and muscle-like populations identified through deconvolution, whereas a separate territory was enriched for EMT and TNFα/NF-κB signaling activity (Fig. 5G). Additional enrichment of KRAS-associated signaling was also observed within good-outcome tissue.

Although the spatial cohort was limited in size and these observations should therefore be interpreted with appropriate caution, the patterns were remarkably consistent across spatial resolutions and independent transcriptomic modalities. Most importantly, the spatial analyses demonstrated that the proliferative programs associated with poor clinical outcome are concentrated within discrete tissue territories, forming localized proliferative niches rather than diffuse transcriptional programs distributed throughout the tumor.

Collectively, these findings suggest that aggressive ACC is characterized not only by expansion of proliferative luminal tumor populations but also by their spatial organization into MYC/E2F-enriched proliferative niches. In contrast, favorable-outcome tumors retain a more heterogeneous architecture composed of differentiated epithelial, stromal, and contractile programs. These observations provide spatial validation of the lineage-proliferation axis identified throughout this study and support the concept that aggressive ACC is driven by a coordinated biologic state spanning cellular composition, transcriptional activity, and tissue organization.

### A Four-Gene Prognostic Signature Captures the Proliferative Luminal State Associated with Aggressive ACC

The remarkable consistency of findings across the single-cell, bulk transcriptomic, coexpression network, and spatial analyses suggested that poor clinical outcome in ACC is driven by a common underlying biological program. We therefore asked whether this biology could be distilled into a useful prognostic classifier. To address this, we developed a three-level hierarchical Bayesian shrinkage model that integrated evidence from differential expression analysis, coexpression networks, and single-cell transcriptomics, and trained it exclusively on our (internal) bulk RNA-seq discovery cohort.

Among 12,700 evaluable genes, the model converged on a four-gene signature consisting of DSCAM, ODC1, NCAPG, and CCNB2, all of which were positively associated with poor clinical outcome. The resulting locked score was defined as the unweighted mean of rank inverse-normal transformed expression values for the four genes, preserving the directionality of each coefficient. Internal cross-validation of the discovery cohort yielded an area under the receiver operating characteristic curve (AUC) of 0.783 ± 0.021 (Fig. S7A).

We next evaluated the locked classifier in independent external cohorts without retraining or recalibration. In the Danish validation cohort (37) (n = 54), which included 21 deaths within five years, the four-gene score significantly predicted overall survival and achieved a concordance index of 0.671 (95% CI, 0.541–0.801) (Fig. 6A-B). Each standard-deviation increase in the score was associated with a 1.74-fold increase in mortality risk (95% CI, 1.10–2.74), and the association remained significant after adjustment for stage and solid histology. Using a threshold determined exclusively from the discovery cohort, patients could be stratified into high- and low-risk groups with significantly different survival outcomes (hazard ratio, 3.38; 95% CI, 1.43–7.98; log-rank P = 0.0033) (Fig. 6A). Importantly, the four-gene classifier outperformed previously published 14-gene and 49-gene ACC signatures evaluated in the same cohort (Fig. 6B).

**Fig. 6.**
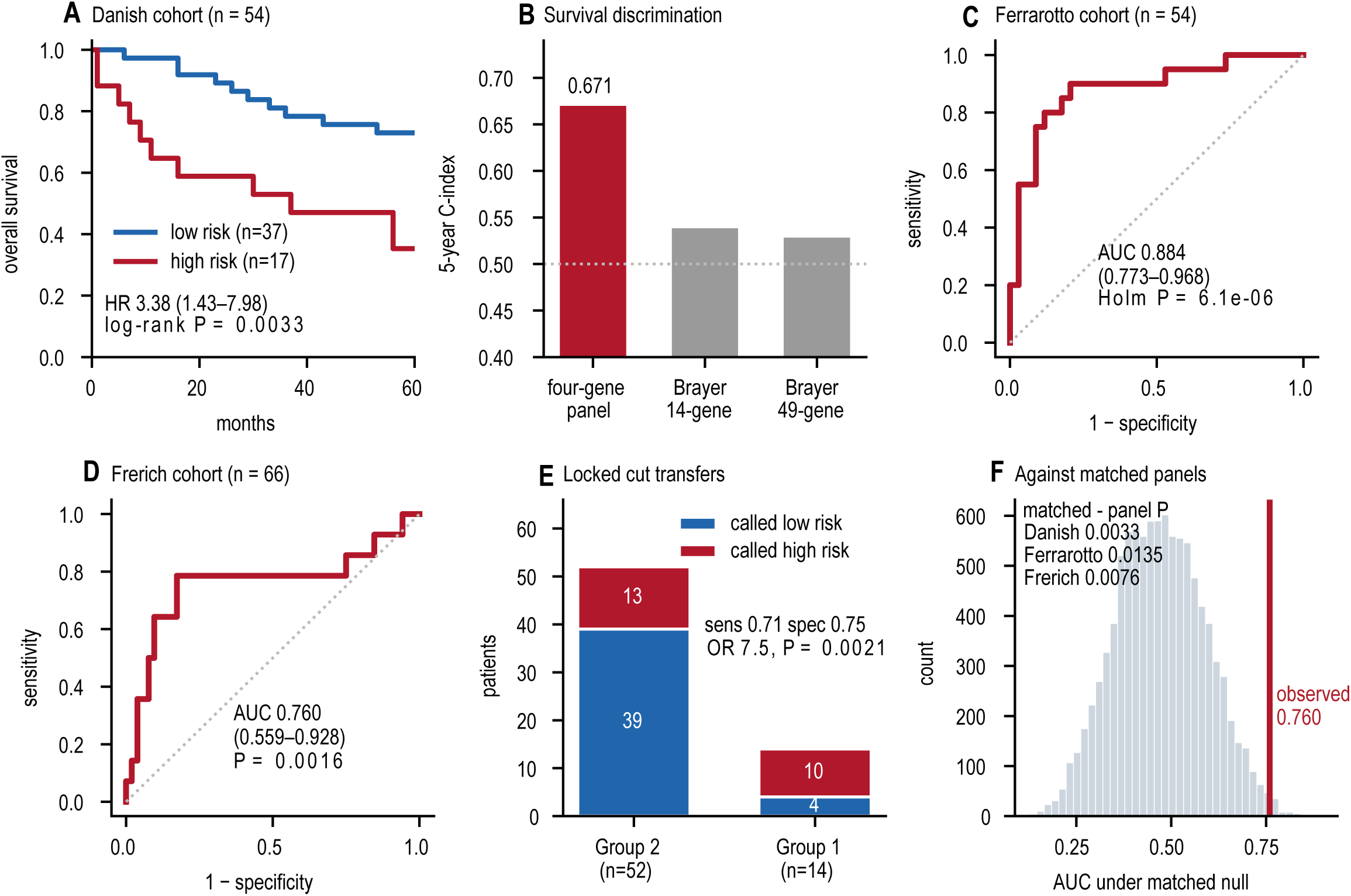
A locked four-gene prognostic classifier captures aggressive ACC biology and validates across independent external cohorts. (A) Kaplan-Meier overall survival analysis of the 54-patient Danish (DK) validation cohort using the locked four-gene classifier. Patients were stratified using a decision threshold derived exclusively from the 20-sample discovery cohort and applied to the validation cohort without retraining or recalibration. (B) Five-year prognostic performance in the Danish cohort measured by Harrell’s concordance index (C-index). The four-gene classifier (DSCAM, ODC1, NCAPG, and CCNB2) is compared with previously published 14-gene and 49-gene ACC prognostic signatures. Statistical comparisons were performed using DeLong’s method for correlated receiver operating characteristic curves. (C) External molecular validation in the Ferrarotto cohort. The four-gene classifier distinguishes the aggressive MYC-driven ACC-I subtype from ACC-II tumors, demonstrating that the score captures biologically meaningful disease states in addition to survival outcomes. (D) Receiver operating characteristic analysis of the Frerich cohort demonstrating identification of clinically aggressive tumors within an independent external dataset. (E) Risk stratification of the Frerich cohort using the same threshold derived exclusively from the discovery cohort. Group 1 represents the poor-outcome/high-risk subgroup, whereas Group 2 includes the remaining tumors. (F) Empirical null benchmarking of classifier performance using 10,000 abundance- and variance-matched random four-gene panels. The distribution of random panel AUC values is shown with the observed performance of the locked classifier indicated. Empirical P values represent the fraction of matched random panels achieving equal or greater performance across the three external validation analyses.

To determine whether the classifier captured biologically meaningful ACC states rather than survival alone, we applied it to additional external datasets. In the Ferrarotto cohort (13) (n = 54), the scores robustly distinguished the aggressive MYC-driven ACC-I subtype from ACC-II tumors, achieving an AUC of 0.884 (95% CI, 0.773–0.968; adjusted P = 6.1 × 10⁻⁶) (Fig. 6C). In the Frerich cohort (38) (n = 66), which included 14 clinically aggressive tumors, the score achieved an AUC of 0.760 (95% CI, 0.559–0.928; P = 0.0016) and correctly classified 10 of 14 high-risk cases (Fig. 6D-E). These findings indicate that the classifier identifies aggressive disease across both survival-based and molecularly defined validation cohorts.

To evaluate whether this performance could be attributed to nonspecific prognostic information, we compared the model against 10,000 abundance- and variance-matched random four-gene panels. Matched random signatures rarely achieved equivalent or superior performance, occurring in only 0.33%, 1.35%, and 0.76% of simulations across the three validation settings (Fig. 6F). These findings indicate that the four-gene panel’s performance substantially exceeds what would be expected by chance.

Interestingly, stability analyses suggested that the biological process captured by the model was more reproducible than the individual genes themselves. Across 45 independent refits generated through alternative random seeds and resampling strategies, no individual panel gene was selected in more than 27% of runs. In contrast, genes associated with G2/M cell-cycle regulation were selected in 56% of refitted models compared with only 1.7% expected by chance (Fig. S7B). Despite differences in gene composition, prognostic performance remained largely stable across refitted models, and even signatures sharing no genes with the locked panel generated positively correlated risk scores (Fig. S7C-D). Similar patterns were observed in both external validation cohorts. These findings suggest that the model captures a robust proliferative program rather than dependence on a specific set of marker genes.

Finally, we examined whether the prognostic signal reflected the cellular states identified throughout this study. Using the internal single-cell atlas as a reference, DWLS deconvolution was applied to the Danish validation cohort. Tumors classified as high risk showed increased abundance of ADCs (Dividing) and LETCs (Epithelial), whereas low-risk tumors were enriched for myoepithelial tumor cells (Myoepithel) (Fig. S7E). The degree of cellular-state enrichment closely tracked the prognostic effects observed in the validation cohort (ρ = 0.80), indicating that the classifier captures the same lineage-proliferation axis identified through the single-cell analyses.

Taken together, these findings demonstrate that a compact four-gene signature derived from integrated multi-omic analyses captures a biologically coherent proliferative program that generalizes across independent patient cohorts. More broadly, they indicate that prognosis in ACC is governed less by individual genes than by a conserved proliferative luminal state characterized by expansion of actively dividing tumor cells, activation of cell-cycle programs, and loss of myoepithelial identity. This lineage-proliferation axis emerged consistently across cellular composition, developmental trajectories, transcriptomic analyses, coexpression networks, and spatial organization, supporting its role as a fundamental determinant of aggressive ACC biology.

### Immunofluorescence of Matched Tumor Sections Reiterate Spatial Expression Patterns at the Protein Level

The spatial analyses placed the proliferative program of poor-outcome tumors within discrete tumor nests, but they measured RNA rather than protein. To test whether the corresponding proteins showed the same distribution, we stained serial sections from the two tumors profiled by Visium HD, the favorable-outcome tumor A742434 and the poor-outcome tumor A368571 (Table 1) immunofluorescence (IF). The IF panel included interesting genes highlighted in the study such MYB, MYC, NOTCH1, ODC1 and SEMA3A for staining. (Fig. 7; Fig. S8).

**Fig. 7.**
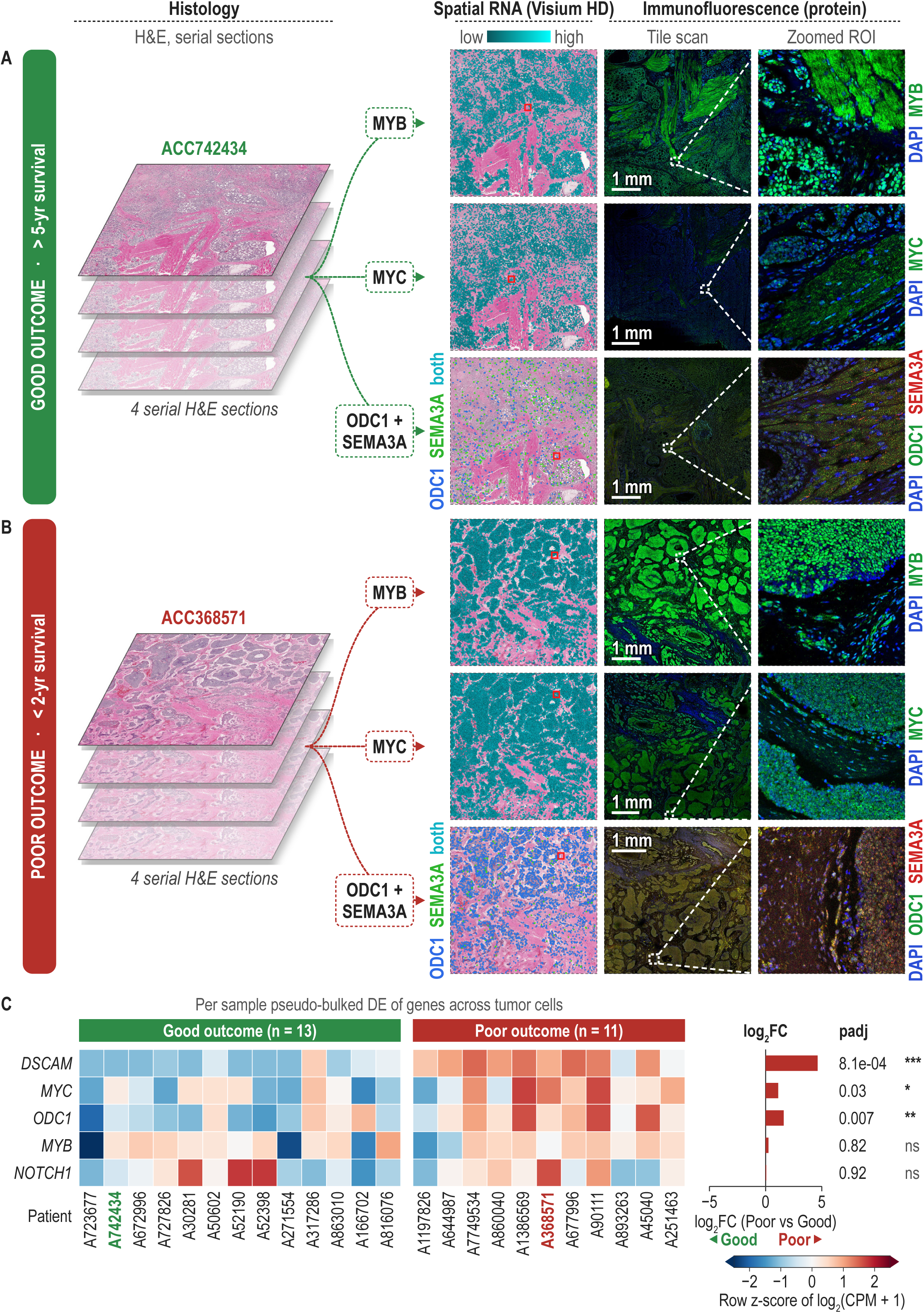
Immunofluorescence of matched Visium HD tumors recapitulates spatial expression patterns at the protein level. (A) Favorable-outcome tumor A742434 (overall survival >5 years). (B) Poor-outcome tumor A368571 (overall survival <2 years). For each tumor, the left column shows serial H&E-stained FFPE sections from the block used for Visium HD profiling. The second column shows Visium HD expression of MYB, MYC, and ODC1 with SEMA3A projected onto the H&E image. MYB and MYC are colored from low to high expression; in the ODC1 and SEMA3A row, bins are colored by detection of ODC1 (blue), SEMA3A (green), or both (cyan). The third column shows whole-section immunofluorescence (IF) tile scans, and the right column shows the regions of interest outlined by boxes at higher magnification. MYB, MYC, and ODC1 are shown in green, SEMA3A in red, and DAPI-stained nuclei in blue. Scale bars, 1 mm (tile scans). (C) Per-patient pseudobulk expression of DSCAM, MYC, ODC1, MYB, and NOTCH1 in tumor cells from the 24 single-cell tumors (13 favorable-outcome and 11 poor-outcome). Heatmap values are row z-scores of log2(CPM + 1). Bars show the PyDESeq2 log2 fold change (poor versus good outcome) with Benjamini-Hochberg-adjusted P values (padj). *padj < 0.05; **padj < 0.01; ***padj < 0.001; ns, not significant.

Protein staining followed the Visium HD expression maps closely (Fig. 7A, B). In the poor-outcome tumor, MYB and MYC transcripts were concentrated in large, densely packed tumor nests, and IF showed strong MYB and MYC staining in the same nests. In the favorable-outcome tumor, both transcripts were confined to smaller, scattered tumor regions within a heterogeneous stroma, and protein staining was correspondingly sparser. MYB-positive tumor cells were present in both specimens, and MYB expression in tumor cells did not differ between outcome groups (FDR = 0.82; Fig. 7C). MYC behaved differently. Its tumor-cell expression was higher in poor-outcome patients (adjusted P = 0.03; Fig 7C), and MYC staining was weak across most of the favorable-outcome section.

ODC1 and SEMA3A were stained together on one section from each tumor. By IF, ODC1 staining was stronger and more widespread among tumor cells of the poor-outcome specimen, in line with its higher tumor-cell expression in poor-outcome patients (adjusted P = 0.007; Fig. 7A, B; Fig. 7C). SEMA3A was more detected in scattered cells of the favorable-outcome tumor, consistent with the enrichment of SEMA3A-NRP1 signaling in favorable-outcome tumors.

NOTCH1 RNA transcripts and NOTCH1 staining on tissue were both concentrated in tumor nests of the poor-outcome specimen, whereas NOTCH1 protein was barely detectable in the favorable-outcome tumor (Fig. S8 A, B). NOTCH1 RNA transcripts did not differ between outcome groups in tumor-cell pseudo bulk profiles (adjusted P = 0.92) (Fig. 7C). The stronger NOTCH1 protein signal in the poor-outcome tumor may therefore reflect post-transcriptional regulation or the tumor-cell states, which agrees with the higher NOTCH pathway activity measured in malignant cells through pseudotime (Fig. S3F). Taken together, the matched IF and Visium HD data at the same location shows agreement between the RNA expression and the protein levels in the tumor tissue substantiating our Visium spatial analysis.

## Discussion

Head and neck adenoid cystic carcinoma remains a clinically challenging malignancy because patients with apparently similar clinicopathologic features frequently experience markedly different outcomes, yet reliable molecular biomarkers capable of identifying high-risk disease at diagnosis remain lacking. By integrating bulk RNA sequencing, single-cell RNA sequencing, and spatial transcriptomics, we identified and externally validated a reproducible outcome-associated lineage-proliferation axis in HNACC. Poor-outcome tumors were characterized by profound depletion of myoepithelial tumor cells, expansion of luminal epithelial and actively dividing tumor populations, activation of MYC-, E2F-, and G2/M-associated programs, and emergence of proliferative spatial niches. In contrast, favorable-outcome tumors retained myoepithelial, contractile, and extracellular matrix-associated states. These patterns emerged consistently across orthogonal analytical approaches, suggesting that clinical outcome in HNACC is linked to a transition from a myoepithelial-rich state toward a proliferative luminal state.

The most important finding of this study is the identification of a clinically relevant myoepithelial-to-luminal transition associated with outcome. ACC has long been recognized as a biphasic tumor composed of luminal epithelial and myoepithelial compartments, and previous transcriptomic studies suggested that myoepithelial-rich tumors display less aggressive behavior (39). Single-cell profiling further showed substantial variation in luminal and myoepithelial content and identified a paracrine interaction in which myoepithelial cells supply NOTCH ligands to neighboring luminal populations (21). Our findings place these observations into a clinical context. Myoepithelial tumor cells declined from approximately 50% of malignant cells in favorable-outcome tumors to less than 4% in poor-outcome tumors, whereas luminal epithelial populations expanded more than threefold. Importantly, the increase in actively dividing tumor cells was driven largely by expansion of the luminal lineage itself rather than increased proliferation within individual luminal cells. This distinction could not have been resolved using bulk transcriptomics alone and highlights the value of integrating cellular-resolution approaches to study rare cancers.

The biological implications of myoepithelial depletion likely extend beyond cell-type abundance. Myoepithelial cells contribute to the maintenance of extracellular matrix organization, tissue structure, and epithelial homeostasis. Loss of myoepithelial integrity has similarly been associated with progression and invasion in other biphasic epithelial malignancies (40). Our data suggest that a comparable process may occur in HNACC, whereby progressive loss of the myoepithelial compartment permits luminal expansion and facilitates acquisition of proliferative programs. Consistent with this interpretation, favorable-outcome tumors retained extracellular matrix organization, myogenesis, smooth-muscle contraction, and contractile signatures across bulk, single-cell, and spatial datasets.

Trajectory analyses further refined this model. Monocle3 and CytoTRACE2 consistently positioned myoepithelial-like tumor cells near the root of the inferred developmental continuum and identified progressively increasing luminal differentiation along pseudotime. Although favorable- and poor-outcome tumors shared a common developmental architecture, their terminal states diverged substantially. Poor-outcome tumors converged upon a highly proliferative endpoint dominated by actively dividing tumor cells, whereas favorable-outcome tumors retained more differentiated luminal populations. These findings suggest that aggressive ACC does not arise from a distinct lineage but rather reflects progression along a common developmental trajectory toward a proliferative luminal endpoint. This interpretation is consistent with previous reports demonstrating luminal enrichment and increased NOTCH activity in recurrent and metastatic ACC (11,21).

Our study also adds context to the role of NOTCH signaling in ACC progression. Activating NOTCH1 alterations are among the best-established markers of aggressive disease and have been linked to recurrence, metastasis, and poor prognosis (11–13). In the present study, NOTCH-associated transcriptional programs increased progressively along pseudotime and were consistently elevated within malignant cells from poor-outcome tumors. At first glance, this appears discordant with our ligand-receptor analyses, where DLL1-NOTCH1 interactions were more prominent in favorable-outcome tumors. However, ligand-receptor analyses evaluate one potential upstream mechanism of pathway activation, whereas pathway scores quantify integrated downstream transcriptional output. Thus, elevated NOTCH activity in aggressive tumors can coexist with preservation of individual ligand-receptor interactions in more differentiated tumors. Together, these findings support a model in which NOTCH signaling is embedded within a broader differentiation and proliferation continuum rather than defining a discrete malignant population.

Across every transcriptomic modality examined, proliferation emerged as the dominant molecular feature of poor clinical outcome. Bulk RNA sequencing identified enrichment of MYC targets, E2F targets, G2/M checkpoint pathways, chromosome segregation, DNA replication, ribosome biogenesis, translation, and RNA-processing programs. Similar signatures were independently recovered through pseudobulk analyses of luminal tumor populations and through outcome-associated WGCNA modules. These findings closely parallel the ACC-I subtype described by Ferrarotto and colleagues, which is characterized by MYC activation, NOTCH signaling, and poor prognosis (13). These findings are also consistent with prior transcriptomic studies describing stem-like and proliferative ACC subgroups associated with adverse outcomes (38). The remarkable consistency of these signals across independent analytical frameworks strongly suggests that MYC/E2F-associated cell-cycle activation is a fundamental feature of aggressive ACC biology.

Our findings also raise the question of whether the proliferative luminal population identified in HNACC shares features with luminal states observed in other epithelial malignancies. In breast cancer and other glandular tumors, luminal differentiation programs are frequently linked to enhanced proliferative capacity and MYC/E2F activation (41). While HNACC is biologically distinct and should not be considered directly analogous to breast cancer, the convergence on luminal identity together with proliferation-associated programs raises the possibility that common lineage-associated mechanisms contribute to aggressive behavior across multiple epithelial tumor types. Whether these proliferative luminal states share therapeutic vulnerabilities remains an important area for future study.

Interestingly, the favorable-outcome state appeared to represent more than simply the absence of proliferation. In addition to retaining myoepithelial populations, favorable-outcome tumors exhibited enrichment of WNT5B-associated networks, extracellular matrix organization, contractile programs, and tissue differentiation signatures. WNT5B emerged as a hub gene within the favorable-outcome coexpression module and has been linked to noncanonical WNT signaling, tissue organization, and maintenance of differentiated cellular states in multiple biological contexts (42). Although our data do not directly demonstrate a dormant or stem-like state, the preservation of differentiated stromal and myoepithelial programs, together with limited proliferative activity, raises the possibility that long-term surviving tumors occupy a more constrained cellular state that slows progression despite the presence of established oncogenic drivers.

Spatial transcriptomics provided orthogonal validation while preserving tissue architecture. Poor-outcome tumors exhibited tumor-rich territories enriched for actively dividing tumor cells and MYC/E2F-associated programs, whereas favorable-outcome tumors displayed greater cellular and transcriptional heterogeneity, including mural, epithelial, and extracellular matrix-associated territories. Importantly, proliferative programs were not uniformly distributed throughout the tissue but instead localized within discrete spatial niches. Immunofluorescence of matched sections showed the same organization at the protein level, with strong MYC and NOTCH1 staining concentrated in the tumor nests of the poor-outcome tumor. These observations suggest that the proliferative luminal state identified by bulk and single-cell analyses is physically organized within the tumor microenvironment and may represent a distinct tumor ecosystem associated with aggressive disease.

Our communication analyses provided additional insight into the biologic differences between outcome groups. Clinically, perineural invasion is common across ACC and can be observed even among patients with relatively favorable outcomes (43). Interestingly, although neural-associated signaling was present in both cohorts, the pathways involved differed. Favorable-outcome tumors demonstrated enrichment of SEMA3A-NRP1 signaling, whereas poor-outcome tumors preferentially utilized NTF3-NTRK3, NTF3-NTRK2, and NTN1-DSCAM interactions. Semaphorin signaling has recognized roles in axonal guidance and neuron-tumor interactions (44,45), suggesting that neurotropism itself may not distinguish favorable-from poor-outcome disease. Instead, distinct neural signaling programs may operate within different biologic contexts. The recurrent appearance of NTRK- and DSCAM-associated interactions in poor-outcome tumors is particularly intriguing given the propensity of aggressive ACC for perineural spread and warrants further investigation.

The four-gene classifier developed in this study provides a potential translational application of these findings. The locked panel comprising DSCAM, ODC1, NCAPG, and CCNB2 predicted survival in an independent Danish cohort and outperformed previously reported ACC classifiers (37) and successfully identified aggressive disease states across additional validation cohorts.

Importantly, model-refitting analyses demonstrated that the underlying biological process was substantially more stable than the individual genes themselves. Multiple refits repeatedly converged on G2/M and proliferation-associated programs despite considerable variation in gene composition, suggesting that the classifier functions primarily as a surrogate for a broader biologic state rather than relying on a small set of uniquely causal transcripts. This likely explains its transferability across independent datasets.

DSCAM warrants particular attention because it emerged repeatedly across differential-expression, communication, and prognostic analyses. Although its role in ACC remains undefined, its established functions in neuronal guidance and neural development make it an attractive candidate linking aggressive disease, neurotrophic signaling, and perineural invasion. Future studies are needed to determine whether DSCAM drives progression or simply serves as a robust marker of aggressive disease states.

Unlike the strong lineage and proliferative signals identified here, immune-cell abundance did not consistently distinguish outcome groups. This observation is consistent with previous reports describing ACC as an immunologically cold tumor with limited immune infiltration and poor responsiveness to immunotherapy (14,17,46). Although several communication pathways were enriched in poor-outcome tumors, patient-level statistical power was limited, and these observations should therefore be regarded as hypothesis-generating.

Several limitations should be acknowledged. Although this represents one of the largest integrated multi-omic studies of HNACC assembled to date, cohort sizes remain constrained by disease rarity. The study was retrospective and will require prospective validation in independently collected patient cohorts. Spatial analyses were performed on a limited number of tumors and should therefore be interpreted as orthogonal validation rather than definitive population-level characterization. Apart from that the transcriptionally similar cell types had to be merged for credible deconvolution results. Finally, while the convergent findings strongly support a myoepithelial-to-proliferative luminal transition, functional studies are needed to determine whether this transition is causally linked to disease progression and whether it represents a therapeutic vulnerability.

In conclusion, our study identifies a reproducible lineage-proliferation axis that links cellular composition, developmental state, transcriptional programming, and tissue architecture to clinical outcome in HNACC. Aggressive tumors are characterized by loss of a myoepithelial-rich state, expansion of proliferative luminal populations, activation of MYC/E2F/G2/M-associated programs, and emergence of spatially organized proliferative niches. The reproducibility of this axis across bulk, single-cell, and spatial transcriptomic platforms, along with external validation, suggests it is a fundamental determinant of ACC progression. These findings provide a biologically grounded framework for prognostic stratification and support future investigation of NOTCH-, neurotrophic-, cell-cycle-, and proliferation-directed therapeutic strategies in high-risk ACC.

## Methods

### Human Samples and Study Design

Tumor specimens from patients with adenoid cystic carcinoma (ACC) were obtained from Dr. El-Naggar at The University of Texas MD Anderson Cancer Center under Institutional Review Board-approved protocols with informed consent. After a material transfer agreement and an Institutional Review Board-approved protocol at the University of New Mexico, we collected and analyzed the tumor samples. Patients were grouped by clinical outcome, with a favorable outcome defined as overall survival of more than 5 years and a poor-outcome as overall survival of less than 2 years. The study was conducted in accordance with the ethical principles of the Declaration of Helsinki.

Patient sex was not collected as part of the clinical metadata for this study; hence, sex was not considered in the study design or analyses.

The bulk RNA Seq cohort comprised 20 head and neck ACC (HNACC) tumors from 10 anatomical sites, including 10 favorable-outcome and 10 poor-outcome tumors. A single-cell RNA-seq cohort comprised 24 tumors (13 favorable-outcome and 11 poor-outcome) from 9 head and neck sites, including 19 tumors overlapping with the bulk RNA-seq cohort. Spatial transcriptomic profiling was performed on six tumors using the 10x Genomics Visium platform, including four standard-resolution Visium samples and two Visium HD samples. Three published bulk RNA-seq cohorts were used only for external validation: a Danish survival cohort (n = 54) (37), the Ferrarotto cohort with ACC-I/ACC-II subtype labels (n = 54) (13) and the Frerich cohort with clinically aggressive tumors annotated (n = 66) (38). Clinical characteristics of tumor samples is summarized in **Table 1**.

### Bulk RNA Sequencing and Transcriptomic Analysis

Bulk RNA sequencing was performed using formalin-fixed, paraffin-embedded (FFPE) tumor specimens. Total RNA was isolated using standard RNA extraction workflows and assessed using an Agilent Bioanalyzer 2100. RNA integrity numbers (RINs) ranged from 2.2 to 2.5, consistent with FFPE-derived material. Each tissue sample was first suspended in RLT lysis buffer and mechanically disrupted with a plastic pestle. After lysis, tissue was homogenized with QIAshredders (Qiagen). RNA was extracted from the resulting lysate using EZ2 RNA/miRNA Tissue/Cell Kits (Qiagen). Sequencing run was performed on the Singular G4 platform with 25B reagent kit (300 Cycle). Illumina BCL Convert v3.9.3 convert base call (BCL) files into FASTQ files.

Libraries were prepared according to standard RNA-seq protocols and sequenced on the Singular Genomics G4 platform using paired-end 150 × 150 bp sequencing. Reads were aligned to the GRCh38 human reference genome using STAR, v2.7.5a (47), to generate gene-level count matrices for downstream analyses. Global transcriptional relationships among tumors were assessed using principal component analysis. Differential expression analyses between favorable- and poor-outcome tumors were performed using PyDESeq2, with design factors taken from the clinical metadata and the contrast computed with the DeseqStats module. Genes with an adjusted P value <0.05 and absolute log2 fold change >0.5 were considered differentially expressed.

Biological pathway activity was evaluated using gene set enrichment analysis (GSEA) and over-representation analysis (ORA) against MSigDB, Reactome, Gene Ontology, and KEGG pathway collections. GSEA was implemented with the prerank function of GSEApy, using genes ranked by the PyDESeq2 test statistic, 1,000 permutations, a random seed of 6 and a minimum gene set size of 15. The 2025 release of the MSigDB, Hallmark, Oncogenic Signatures, KEGG, Reactome and Gene Ontology (Biological Process, Cellular Component and Molecular Function) collections was stored locally to ensure reproducibility. Positive normalized enrichment scores (NES) denote enrichment in the good-outcome cohort and negative NES in the poor-outcome cohort. Weighted gene co-expression network analysis (WGCNA) was used to identify outcome-associated transcriptional modules and hub genes.

WGCNA was performed with the WGCNA R package. Raw counts from the 20 samples were filtered to genes with at least 10 counts in at least 50% of samples and normalized by variance-stabilizing transformation in DESeq2. Sample clustering and goodSamplesGenes inspection identified one outlier (A816076, good outcome), which was removed, and the bottom 25% of genes by variance were excluded. Scale-free topology analysis gave a soft-thresholding power of 14 for the signed, weighted network, and modules were detected with blockwiseModules. Module-trait relationships were estimated by Pearson correlation of module eigengenes with clinical outcome, with significance from Student’s asymptotic P value test. Gene significance was defined as the correlation of gene expression with the outcome trait, module membership (kME) was calculated, and hub genes were the top 20 genes per module ranked by kME. Intramodular connectivity (kWithin) was calculated to characterize gene centrality. Module networks were built with igraph (48), retaining edges with a topological overlap (TOM) of at least 0.225 (blue module) or 0.185 (turquoise module). Over-representation analysis (ORA) of module genes used the enrichr function of GSEApy against the same locally stored MSigDB collections, with the 16,639 genes remaining after WGCNA filtering as the background and a P value cutoff of 0.05.

### Single-Cell RNA Sequencing Analysis

Fresh frozen tumor specimens were dissociated into single-cell suspensions and profiled using the 10x Genomics Chromium Fixed RNA Profiling platform using CG000477 protocol. Libraries were sequenced on the Singular Genomics G4 platform using a 28 × 90 bp read configuration and processed using Cell Ranger (49).

Cell Ranger output was processed in Python with Scanpy (24). Filtering was done independently per sample, retaining cells with at least 200 detected genes and genes expressed in at least 3 cells. Cells with more than 10% mitochondrial counts were excluded as potentially dead cells. Doublets were called with DoubletDetection (50), Scrublet (51) and SOLO (52), and cells flagged by at least two of the three methods were removed. As a final filter, cells above the 95th percentile of the total count distribution were excluded as potential doublets or outliers. Cells were then normalized and log1p transformed, the top 10,000 highly variable genes were selected with the seurat_v3 flavor in Scanpy, and S and G2/M cell-cycle scores were computed and regressed out to limit confounding by proliferation state. Following quality control, datasets were integrated using single-cell variational inference (scVI) (25) to mitigate batch effects. The scVI model used a 30-dimensional latent space and was trained for 300 epochs with validation checkpoints every 10 epochs; its latent representation was used for the neighborhood graph, UMAP and Leiden clustering at a resolution of 0.6. Cell populations were identified through unsupervised clustering and annotation using established marker genes.

### Pseudobulk Differential Expression

Sample-level differential expression was performed on pseudobulk profiles, summing counts across all cells of the relevant population in each sample, with PyDESeq2 (31) and a design based on clinical outcome (13 good and 11 poor). Significant genes were called at an adjusted P value < 0.05 and absolute log2 fold change > 0.5. Sensitivity analyses were performed using repeated cell down-sampling and count-thinning procedures to evaluate robustness of pseudobulk differential-expression findings.

### Cell-Type Compositional Analysis

Differential cell-type abundance between outcome groups was evaluated using scCODA (v0.1.9) (26). Single-cell annotations were aggregated into a sample-by-cell-type count matrix, with outcome as a categorical covariate, good outcome as the reference level and mural cells as the reference cell type. The model was fitted by Hamiltonian Monte Carlo sampling with 100,000 posterior samples after 10,000 burn-in iterations and fixed random seeds. Effects were assessed by posterior inclusion probability and scCODA’s data-adaptive threshold targeting an estimated false discovery rate (FDR) < 0.05, with 94% highest density intervals extracted from the posterior. Robustness to the reference choice was tested by repeating the analysis with endothelial cells, M2 macrophages and fibroblasts as alternative references. Sample-level cell-type proportions were compared descriptively with two-sided Mann-Whitney U tests.

### Cell-Cell Communication Analysis

Cell-cell communication was inferred using CellPhoneDB (v5.0.1) (27,28) with the statistical analysis method and interaction scoring enabled. Raw counts were normalized to 10,000 transcripts per cell and log1p transformed, and statistical testing used 1,500 permutations with a significance threshold of P < 0.05. A global descriptive interaction map was generated from all samples combined. For outcome-associated analyses, CellPhoneDB was run independently for each patient, excluding cell types represented by fewer than 10 cells. A directed endpoint was defined as a unique ligand-receptor interaction and sender-receiver cell-type pair and was considered evaluable for a patient only when both cell types had at least 10 cells and CellPhoneDB returned a finite, significant interaction score.

Per-patient interaction scores were compared between outcome groups for endpoints evaluable in at least six patients per group, using two-sided Mann-Whitney U tests (exact tests for tie-free distributions and the asymptotic test with tie correction otherwise). Effect sizes were rank-biserial correlations, with positive values denoting higher scores in the good-outcome cohort, reported with median differences and 95% percentile confidence intervals from 10,000 stratified bootstrap resamples in which patients were resampled within outcome groups. Benjamini-Hochberg correction was applied across all evaluable endpoints (FDR < 0.05). Fisher’s exact tests, independently Benjamini-Hochberg adjusted, flagged endpoints whose evaluability differed between outcome groups.

In a complementary exploratory analysis, cells were pooled within each outcome group (cell types with more than 10 cells per group), downsampled to a common depth of 42,214 cells per group (seed 0, repeated with seeds 1-4) and compared across shared cell-type pairs of each ligand-receptor interaction with a paired Wilcoxon signed-rank test (FDR < 0.05). Because the unit of replication was cell interactions rather than patients, these results were considered exploratory. Pooled interactions meeting the FDR threshold were classified as directionally consistent when the group with the higher pooled score also showed equal or greater per-patient presence and a higher mean per-patient interaction score.

### Developmental Potential and Trajectory Analysis

Cells labeled as malignant or carrying a tumor tag were retained, giving 71,984 cells from 24 patients. CytoTRACE2 (29) was applied to raw expression and gene counts, with potency scores calculated in the malignant compartment and then in all annotated cells, and Pearson correlation was used to test the association of CytoTRACE2 scores with S and G2/M phase scores. Trajectory ordering was performed with Monocle3 (30) on the same patient-aligned embedding, with the root selected blind to clinical outcome; two further independent roots were used to assess ordering stability. Program scores (myoepithelial-like, ductal/NOTCH-associated, NOTCH receptor, NOTCH target and EMT) were calculated as the mean standardized expression of the genes in each signature. Per-patient median and mean CytoTRACE2 scores and the fractions of cells above the cohort-wide top 10%, 25% and 33% thresholds (samples with at least 10 malignant cells) were compared between outcome groups using Wilcoxon rank-sum tests and Cliff’s delta effect sizes.

### Spatial Transcriptomic Analysis

Spatial transcriptomic profiling was performed on FFPE tissue sections using the 10x Genomics Visium Spatial Gene Expression and Visium HD platforms with the Human Transcriptome Probe Set v2.0. Standard Visium libraries were sequenced on the Singular Genomics G4 platform using a 28 × 50 bp read configuration, whereas Visium HD libraries were sequenced using a 43 × 50 bp read configuration.

Sequencing data were processed using Space Ranger and analyzed using Giotto (v4.4.3) (34). Each sample was processed independently in Giotto, with Visium HD data (2 × 2 µm bins) aggregated to 16 × 16 µm bins. Filtered feature-barcode matrices and spatial coordinates were loaded with createGiottoVisiumObject and restricted to spots or bins under tissue. Mitochondrial and ribosomal genes were removed before quality-control filtering. Bins were retained if at least 50 (16-µm bins) or 200 (55-µm spots) genes were detected, and genes were retained if detected in at least 50 bins (Visium HD) or 10 spots (standard Visium). Counts were normalized and log-transformed with normalizeGiotto. Highly variable genes were identified with calculateHVF, the first 15 principal components were used for UMAP and shared-nearest-neighbor graph construction (k = 15), and unsupervised clustering used the Leiden algorithm at a resolution of 0.5. A Delaunay triangulation spatial network (minimum k = 2) was used for all downstream spatial analyses. Spatially variable genes were identified with binSpect (rank-based binarization) with Benjamini-Hochberg correction, and the 250 most significant genes were used to detect spatial co-expression modules (metagenes) with detectSpatialCorFeats, partitioned by Ward-linkage hierarchical clustering of gene-gene correlation distances, with the number of modules (k = 2-12) chosen per sample by mean silhouette width and within-cluster sum of squares.

Cell-type compositions were inferred using SpatialDWLS deconvolution (35) with the scRNA-seq atlas generated in this study as the reference. Metagene modules were tested against the MSigDB Hallmark, Gene Ontology Biological Process and Molecular Function, and Reactome collections (release 2026.1.Hs). Over-representation used the exact hypergeometric test against the background of genes passing quality control, with Benjamini-Hochberg correction. Gene-set enrichment used fgsea (53) (minSize = 10, maxSize = 500, nPermSimple = 10,000) on genes ranked by Pearson correlation with each module’s metagene score, excluding the module’s own member genes from the primary ranking. For deconvolution, compositions were summarized per sample for each Leiden domain and for spatial blocks of 400 × 400 µm (Visium HD) or a 600 × 600 µm hexagonal lattice (standard Visium). Immune infiltration was summarized as the summed, unnormalized proportion of myeloid, T and B/plasma cells. Cross-sample comparisons between outcome groups were descriptive, and no statistical tests were applied.

### Immunofluorescence Staining and Imaging

IF was performed on FFPE sections of the Visium HD-profiled tumors A742434 and A368571; ODC1 and SEMA3A were co-stained, and MYB, MYC and NOTCH1 were stained on separate serial sections. Slides were baked at 75 °C for 45 min, deparaffinized in xylene (2 × 5 min) and rehydrated through graded ethanol (100%, 95%, 80%, 50% and 30%, 5 min each) to distilled water. For antigen retrieval, slides were heated at 95–100 °C for 20 min in 10 mM sodium citrate (pH 6.0), cooled for 20–30 min and washed in Tris-buffered saline (TBS) with 0.05% Tween-20 (TBST; 3 × 5 min). Sections were blocked with 5% bovine serum albumin (BSA) in TBS with 0.05% Tween-20 (TBST) for 30–60 min at room temperature.

Primary antibodies (Supplementary Table 1) were diluted in antibody diluent containing 1% BSA and applied (50–100 µl per section) overnight at 4 °C in a dark, humidified chamber. After TBST washes with gentle agitation (3 × 5 min), sections were incubated with fluorophore-conjugated secondary antibodies (Supplementary Table 1) diluted in blocking buffer for 1–2 h at room temperature in the dark, washed in TBST (3 × 5 min), counterstained with DAPI for 5 min and briefly rinsed. Autofluorescence was then quenched with TrueBlack (from Biotium, cat. no. 23011) for 30 s, followed by washes in detergent-free TBS (detergent removes TrueBlack) and mounting in antifade mounting medium (Supplementary Table 1). Whole sections were imaged on a Leica TCS SP8 confocal microscope (Leica Microsystems) using spiral tile scanning at 20x objective, and images were deconvolved with Huygens software (Scientific Volume Imaging).

### Prognostic Model Development and Validation

Gene selection and model fitting used only the internal bulk RNA-seq discovery cohort (n = 20) and its binary outcome. Candidate genes were nominated by integrating bulk differential expression, single-cell pseudobulk differential expression in differentially abundant cell types, and WGCNA hub genes from outcome-associated modules, and were retained if supported by at least two of the three sources. Gene selection and effect direction were determined by a three-level hierarchical Bayesian shrinkage model fitted by automatic differentiation variational inference (ADVI) (54).The panel size k was chosen by a parsimony rule applied to internal resampling (leave-one-out across three seeds and repeated five-fold cross-validation), taking the smallest k within 0.02 AUC of the best-performing size. This gave k = 4, all with positive coefficients (higher score, worse prognosis). Panel size was selected during model development rather than specified in advance. The locked score was the unweighted mean of the rank-based inverse-normal-transformed expression of the four genes. Internal performance was re-monitored for information leakage; several quantities had initially been computed once from the 20-patient data rather than within each fold, and correcting this lowered the internal AUC without affecting external validation. Bootstrap optimism correction (Harrell’s method, 1,000 resamples) (55) was applied to the multivariable model. Model stability was assessed with 45 independent refits (20 random-seed and 25 leave-fold-out refits).

Once locked, the panel was applied once, without refitting, to the Danish cohort. Discrimination was measured using Harrell’s C-index (56) and per-standard-deviation Cox hazard ratios at 5 years and over full follow-up, benchmarked against 10,000 abundance- and variance-matched random gene panels and a 2,000-permutation null distribution. We also tested reference single-sample scoring, which maps each patient onto the discovery cohort’s expression distribution rather than the validation batch, and a decision threshold derived solely from the discovery cohort. The panel was compared with published ACC prognostic classifiers (37,38) in the same patients using DeLong’s method for correlated ROC curves (57). In the Ferrarotto and Frerich cohorts, we tested whether the panel recovered the ACC-I/ACC-II subtypes and clinically aggressive tumors, respectively. Bulk cell-type composition of the Danish cohort was estimated by DWLS deconvolution (36) against the 24-sample single-cell atlas after leave-one-patient-out cross-validation against known single-cell composition.

### Statistical Analysis

All statistical analyses were performed in R and Python. Unless otherwise specified, tests were two-sided, and an adjusted P-value or false discovery rate (FDR) <0.05 was considered statistically significant. Spearman correlation was used to evaluate associations between cell populations and molecular features. For prognostic modeling, survival associations were evaluated using Cox proportional hazards models, Kaplan-Meier analyses, and Harrell’s concordance index. Hazard ratios are reported with 95% confidence intervals.

### Use of Generative AI Tools

Microsoft Copilot and Anthropic Claude were used for language refinement and grammatical editing. Anthropic Claude was used to assist in generating analysis code in Python and R. All generated code was manually reviewed and validated by the authors. The authors reviewed, edited, and verified all AI-assisted text and code and take full responsibility for the content of the manuscript.

## Reporting Summary

Further information on research design is available in the Nature Portfolio Reporting Summary linked to this article.

## Data Availability

Raw and processed bulk RNA-seq, single-cell RNA-seq, and Visium and Visium HD data generated in this study have been deposited in NCBI GEO under accession code (released upon manuscript acceptance). The external validation cohorts were obtained from previously published studies: the Danish cohort (37), the Ferrarotto cohort (13), and the Frerich cohort (38). MSigDB gene sets were obtained from https://www.gsea-msigdb.org. All other data supporting the findings of this study are available from the corresponding author on request.

## Code Availability

Code used for all the analysis across all the modalities is available on GitHub under the link: https://github.com/Gopikrishnan-Bijukumar/Multi-omic-Human-Head-and-Neck-ACC-analysis

## Acknowledgements

We thank Dr. Adel K. El-Naggar (The University of Texas MD Anderson Cancer Center) for providing the tumor specimens and the UNM Human Tissue Repository (HTR) for tissue sectioning and support with Visium and Visium HD processing. Figures and schematics were created using BioRender.

## Funding

This work was supported by National Institutes of Health grant R01DE023222 (to V.P.) and by the University of New Mexico Comprehensive Cancer Center Support Grant NCI P30CA118100 through the Analytical and Translational Genomics Shared Resource.

## Author Contributions

G.B. [Conceptualization, Data curation, Formal analysis, Investigation, Methodology, Software, Validation, Visualization, Writing – original draft, Writing – review & editing]; K.J.B. [Conceptualization, Data curation, Investigation, Methodology, Supervision, Validation, Writing – review & editing]; A.V. [Investigation, Methodology, Supervision, Validation, Writing – review & editing]; E.L.B. [Investigation, Writing – review & editing]; D.Y.L. [Validation, Writing – review & editing]; R.K.O. [Validation, Writing – review & editing]; S.A.N. [Validation, Writing – review & editing]; J.S.E. [Conceptualization, Investigation, Methodology, Resources, Supervision, Validation, Writing – review & editing]; V.P. [Conceptualization, Funding acquisition, Methodology, Project administration, Resources, Supervision, Validation, Writing – original draft, Writing – review & editing]. All authors read and approved the final manuscript.

## Competing Interests

The authors declare no competing interests.

## Notes

### Competing Interest Statement

The authors have declared no competing interest.

