## Supplementary Information for "Multi-Omic and Spatial Profiling Defines Outcome-Associated Tumor States in Head and Neck Adenoid Cystic Carcinoma"

#### Supplementary Figure Legends

##### Fig. S1. Quality control, batch correction, and robustness analyses of single-cell luminal lineage transcriptional programs.

(A) UMAP projection of the quality-controlled single-cell dataset prior to integration. Cells are colored by sample of origin, showing substantial sample-specific clustering consistent with batch-associated variation. (B) UMAP projection following scVI-based integration and batch correction. Cells from different tumors are effectively integrated while preserving biologically meaningful cellular structure. (C) Coarse-grained annotation of the integrated single-cell dataset corresponding to the major cell populations identified in the study. (D) Robustness of luminal lineage pseudobulk differential expression analyses to differences in cellular abundance and sequencing depth. Log2 fold changes from the primary analysis are compared with three sensitivity frameworks repeated across 10 independent random seeds: deep (111 cells × 23 patients), wide (400 cells × 20 patients), and count-thinned (198,905 counts × 23 patients). Genes significant in the primary analysis (n = 151) are highlighted. Gene-level effect estimates remained highly concordant across all sensitivity analyses, indicating that the observed transcriptional differences were not driven by unequal cell representation or sequencing depth. (E-G) Gene set enrichment analysis of pseudobulked luminal epithelial tumor cells (LETCs) and actively dividing tumor cells (ADCs) across the full 24-sample single-cell cohort. Enrichment is shown for(E) Gene Ontology Biological Process,(F) Reactome, and(G) MSigDB Hallmark collections. Poor-outcome tumors remain enriched for cell-cycle, mitotic, chromosome segregation, and MYC/E2F-associated programs, whereas favorable-outcome tumors retain differentiation- and muscle-associated transcriptional signatures.

##### Fig. S2. Global and patient-level analysis of outcome-associated cell-cell communication programs in HNACC.

(A) Global cell-cell communication map generated using CellPhoneDB across all 24 HNACC single-cell datasets. Heatmap values indicate the number of significant ligand-receptor interactions detected between sender and receiver cell populations. Warmer colors indicate cell populations participating in more significant interactions. (B) Concordance between pooled and patient-level interaction analyses for the 123 ligand-receptor interactions identified as differentially enriched between outcome groups. An interaction was considered directionally consistent when the group exhibiting the higher pooled interaction score also demonstrated equal or greater interaction prevalence and a higher mean interaction score across individual patients. (C) Representative patient-level interaction scores for six outcome-associated ligand-receptor interactions. For each interaction, sender and receiver cell populations are indicated. Rank-biserial correlation coefficients ($r_{rb}$), 95% bootstrap confidence intervals, and Benjamini-Hochberg-adjusted P values (q values) are shown. "Missing patients" denote samples in which one or both interacting cell populations were represented by fewer than 10 cells and therefore could not be evaluated. (D) Effect-size estimates for the 20 highest-confidence outcome-associated interactions retained after filtering based on pooled-effect direction, patient-level consistency, down-sampling stability, and concordant effect direction across analytical frameworks. Although none remained significant following multiple-testing correction, these interactions represent the most reproducible candidate communication programs distinguishing favorable- and poor-outcome tumors.

**Fig. S3. Robustness of developmental trajectory inference and pseudotime-associated transcriptional programs in HNACC.**

(A-C) Cell-type composition of the trajectory root neighborhood under independent outcome-blinded rooting strategies. (A) Combined cohort (all malignant cells), (B) favorable-outcome tumors, and (C) poor-outcome tumors. Across all analyses, the inferred trajectory root remained consistently enriched for myoepithelial tumor cells, supporting a shared developmental origin. (D-E) Changes in transcriptional program activity across pseudotime percentiles. Program scores were calculated at the single-cell level and summarized along the developmental trajectory. Curves from favorable- and poor-outcome tumors are overlaid to facilitate comparison of pseudotime-associated transcriptional dynamics. (F) Myoepithelial-like and ductal/NOTCH-associated program scores plotted as a function of pseudotime. Myoepithelial-associated programs progressively decline along the trajectory, whereas ductal and NOTCH-associated programs increase, consistent with differentiation toward a luminal tumor state. (G) Patient-level mean expression of NOTCH receptor and NOTCH target gene programs across malignant cell populations. Point size is proportional to the number of cells contributing to each patient-cell type combination. Only samples containing at least 10 malignant cells were included. Poor-outcome tumors demonstrate increased NOTCH pathway activity across multiple malignant populations. (H) Patient-level expression of epithelial-to-mesenchymal transition (EMT)-associated programs across malignant cell populations. Unlike NOTCH signaling, EMT-related programs were detected in both outcome groups and did not demonstrate a consistent association with clinical outcome.

**Fig. S4. Differential gene expression patterns and additional pathway enrichment analyses of the bulk RNA-seq cohort.**

(A) Heatmap showing row-scaled expression (z-score) of the top 20 genes enriched in favorable-outcome tumors and the top 20 genes enriched in poor-outcome tumors across the 20-bulk RNA-seq samples. Samples are annotated by clinical outcome and anatomical site of origin. Outcome-associated expression patterns are observed across multiple anatomical locations, indicating that survival-associated transcriptional programs are not restricted to a specific tumor site. (B-E) Additional preranked gene set enrichment analyses of the bulk RNA-seq cohort using genes ranked by the PyDESeq2 Wald statistic. Enrichment results are shown for (B) Gene Ontology Biological Process, (C) Gene Ontology Cellular Component, (D) Gene Ontology Molecular Function, and (E) KEGG Medicus pathway collections. Across independent pathway databases, poor-outcome tumors consistently demonstrate enrichment of cell-cycle progression, DNA replication, chromosome segregation, RNA processing, ribosome biogenesis, and translational programs, whereas favorable-outcome tumors are enriched for extracellular matrix organization, differentiation-associated pathways, and tissue-structure programs.

**Fig. S5. Quality-control and network-construction parameters for weighted gene co-expression network analysis (WGCNA).**

(A) Hierarchical clustering dendrogram of the 20-bulk RNA-seq tumors based on variance-stabilized expression values. One sample (A816076; asterisk) was identified as an outlier and excluded prior to network construction. (B) Scale-free topology analysis used to select the soft-thresholding power for network construction. A soft-thresholding power of 14 was chosen to generate a signed weighted co-expression network with scale-free network properties. (C) Distribution of genes assigned to individual co-expression modules identified by WGCNA. Modules were generated using a signed network and signed topological overlap matrix (TOM). The gray module represents genes that could not be confidently assigned to a co-expression module and were excluded from downstream biological interpretation. (D) Hallmark pathway over-representation analysis of the turquoise module, which was positively associated with favorable clinical outcome. Enriched pathways are predominantly related to extracellular matrix organization, tissue structure, stromal biology, and differentiation-associated processes, supporting the link between favorable outcome and preservation of differentiated microenvironmental programs.

**Fig. S6. Reference cell-type composition and functional characterization of spatial metagene programs.**

(A) Cell-type composition of the pooled favorable- and poor-outcome single-cell RNA-seq cohorts following collapse into the cellular categories used for SpatialDWLS deconvolution. This integrated single-cell atlas served as the reference dataset for estimation of spatial cell-type proportions in Visium and Visium HD sections. (B) Gene set enrichment analysis (GSEA) of metagene territories identified within the poor-outcome Visium tumor section. Enrichment was performed against the MSigDB Hallmark collection. Only pathways meeting statistical significance (FDR < 0.05) are shown. Poor-outcome metagene territories are enriched for proliferative and tumor-promoting programs, including MYC-, E2F-, and inflammatory signaling-associated pathways. (C) Gene set enrichment analysis of metagene territories identified in the two favorable-outcome Visium tumor sections. Hallmark pathway enrichment is shown above and Reactome pathway enrichment below for each sample. Favorable-outcome territories are enriched for pathways associated with myogenesis, extracellular matrix organization, tissue differentiation, oxidative metabolism, and stromal organization, supporting preservation of a more differentiated tumor microenvironment.

**Fig. S7. Stability, reproducibility, and external validation of the Bayesian-derived four-gene prognostic classifier.**

(A) Internal performance of the Bayesian shrinkage model evaluated in the discovery cohort. Classifier discrimination was assessed both on the full discovery dataset and under fully nested internal validation procedures. Error bars indicate standard deviation across resampling iterations. Correction for information leakage through nested validation modestly reduced internal AUC estimates while preserving the predictive performance observed in independent external validation cohorts. (B) Frequency of gene and pathway selection across 45 independent model refits, including 20 random-seed refits and 25 leave-fold-out refits. Vertical reference lines indicate the frequency expected by chance. While individual genes showed variable selection frequencies, cell-cycle and G2/M-associated programs were repeatedly selected, indicating convergence on a common biological process rather than dependence on a fixed set of transcripts. (C) Five-year Harrell's concordance index (C-index) of refitted models evaluated in the Danish (DK) validation cohort. Performance of each refitted panel is compared with the locked four-gene classifier and abundance- and variance-matched random gene panels. Most refitted models retained predictive performance exceeding that expected by chance. (D) Rank correlation between risk scores generated by the locked four-gene classifier and 13 independently refitted models that shared no genes with the final panel. Despite complete gene-level divergence, alternative models produced highly concordant patient risk rankings, supporting the existence of a shared underlying biological signal. (E) Association between prognostic risk and outcome-associated cellular states in the Danish validation cohort. Bulk RNA-seq samples were deconvolved using DWLS with the internal 24-sample single-cell atlas as reference. The rank of the cohort-level log hazard ratio is plotted against the rank of cell-type compositional shifts identified by scCODA. High-risk tumors preferentially exhibit expansion of proliferative luminal populations and depletion of myoepithelial tumor cells, recapitulating the cellular-state changes identified in the discovery cohort. (F) Performance of each individual gene from the locked four-gene classifier (DSCAM, ODC1, NCAPG, and CCNB2) in the two external molecular validation cohorts. Percentages beneath each gene indicate the proportion of samples in which expression was undetectable. Although each gene demonstrates varying levels of prognostic or molecular discrimination individually, the combined four-gene score consistently outperforms any single-gene predictor.

**Fig. S8. NOTCH1 immunofluorescence in matched Visium HD tumor region and tumor-cell expression of the immunofluorescence targets across the single-cell cohort.**

(A, B) H&E-stained section, Visium HD expression of NOTCH1 projected onto the H&E image (low to high) red box showing the region of interest (ROI), whole-section NOTCH1 immunofluorescence (IF) tile scan, and ROI (white box). (A) The favorable-outcome tumor A742434 (B) The poor-outcome tumor A368571. NOTCH1 is shown in green and DAPI-stained nuclei in blue. Scale bars, 1 mm (tile scans).

#### Supplementary Figure Panel 1

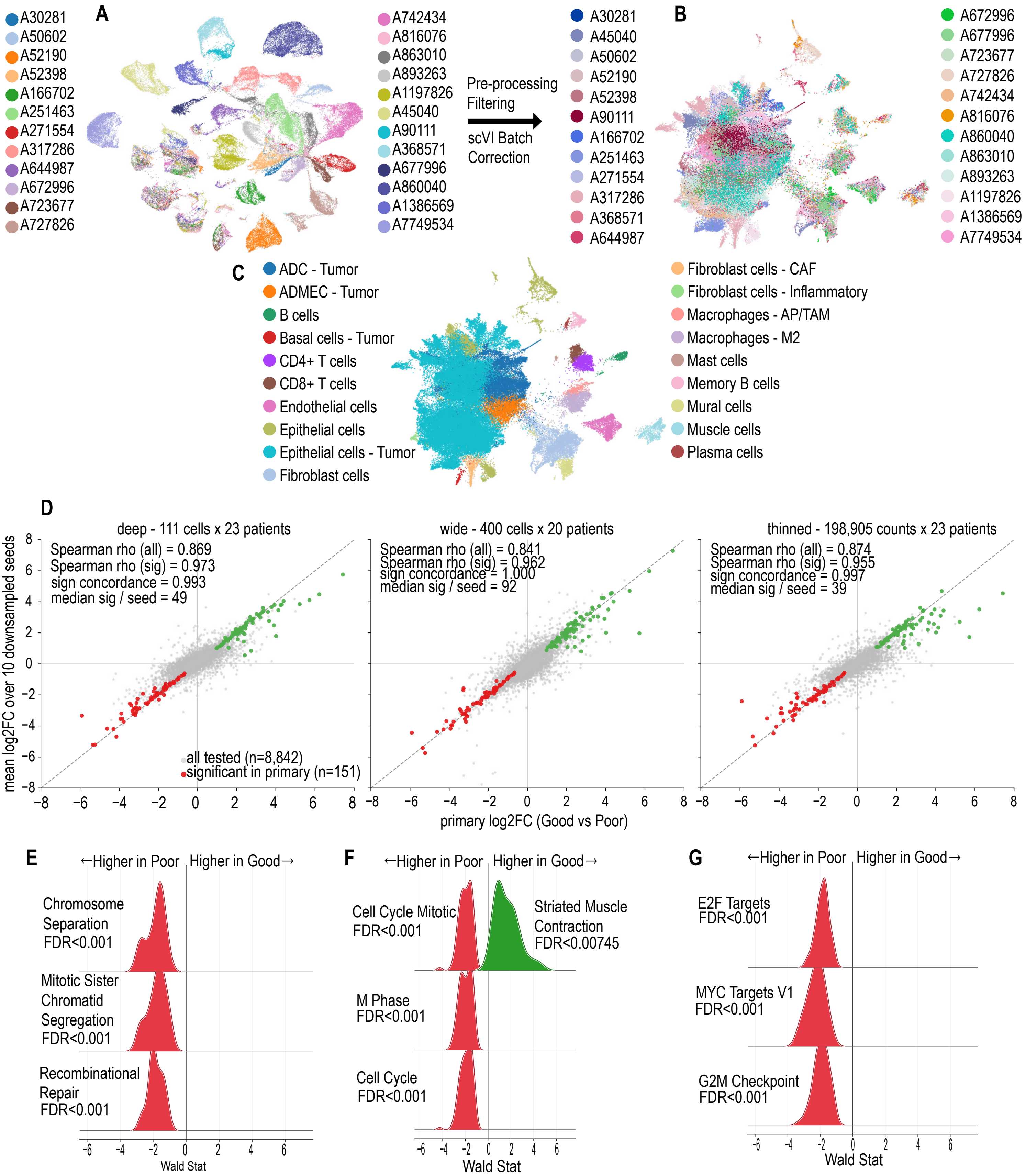

#### Supplementary Figure Panel 2

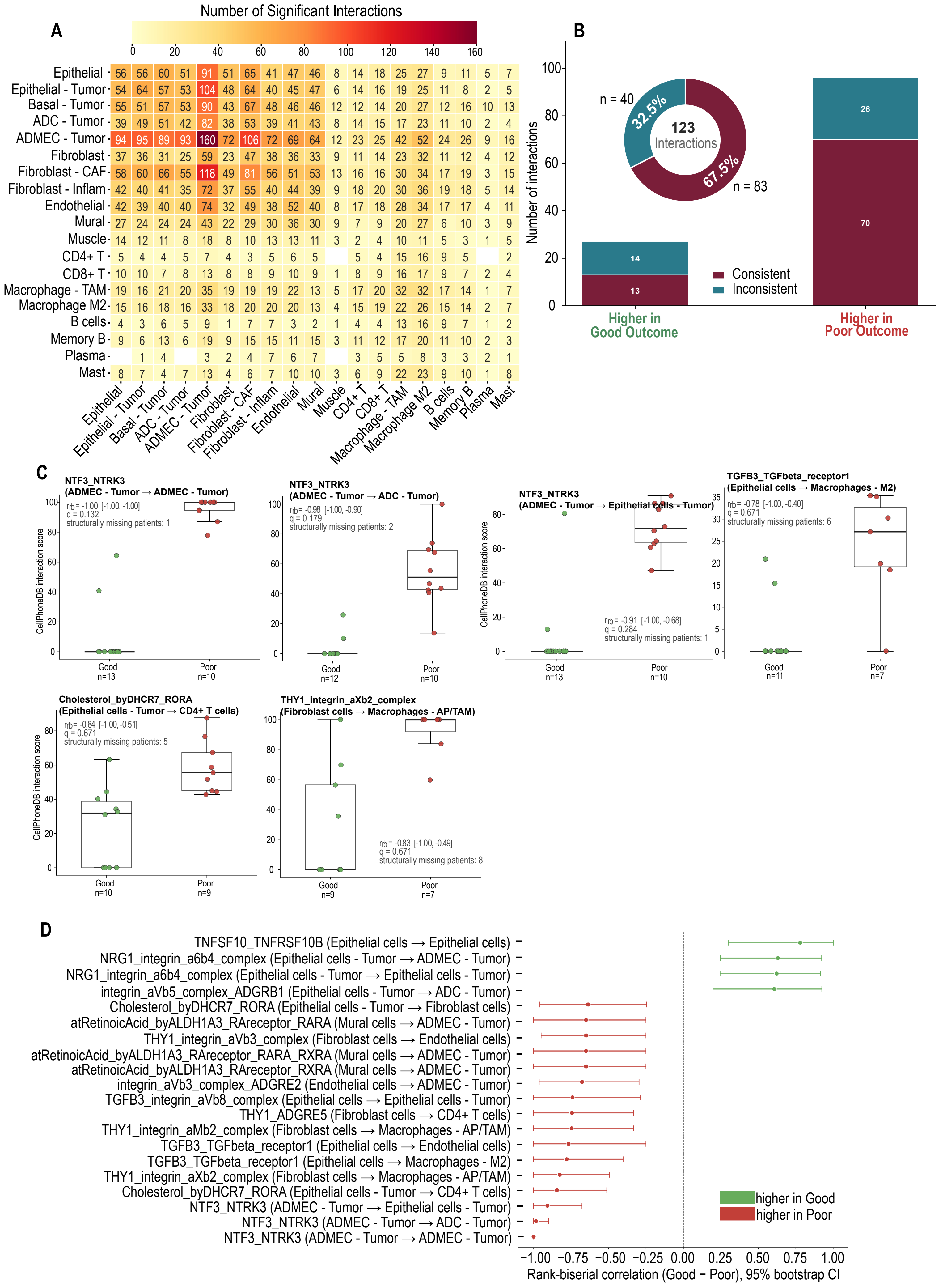

### Supplementary Figure Panel 3

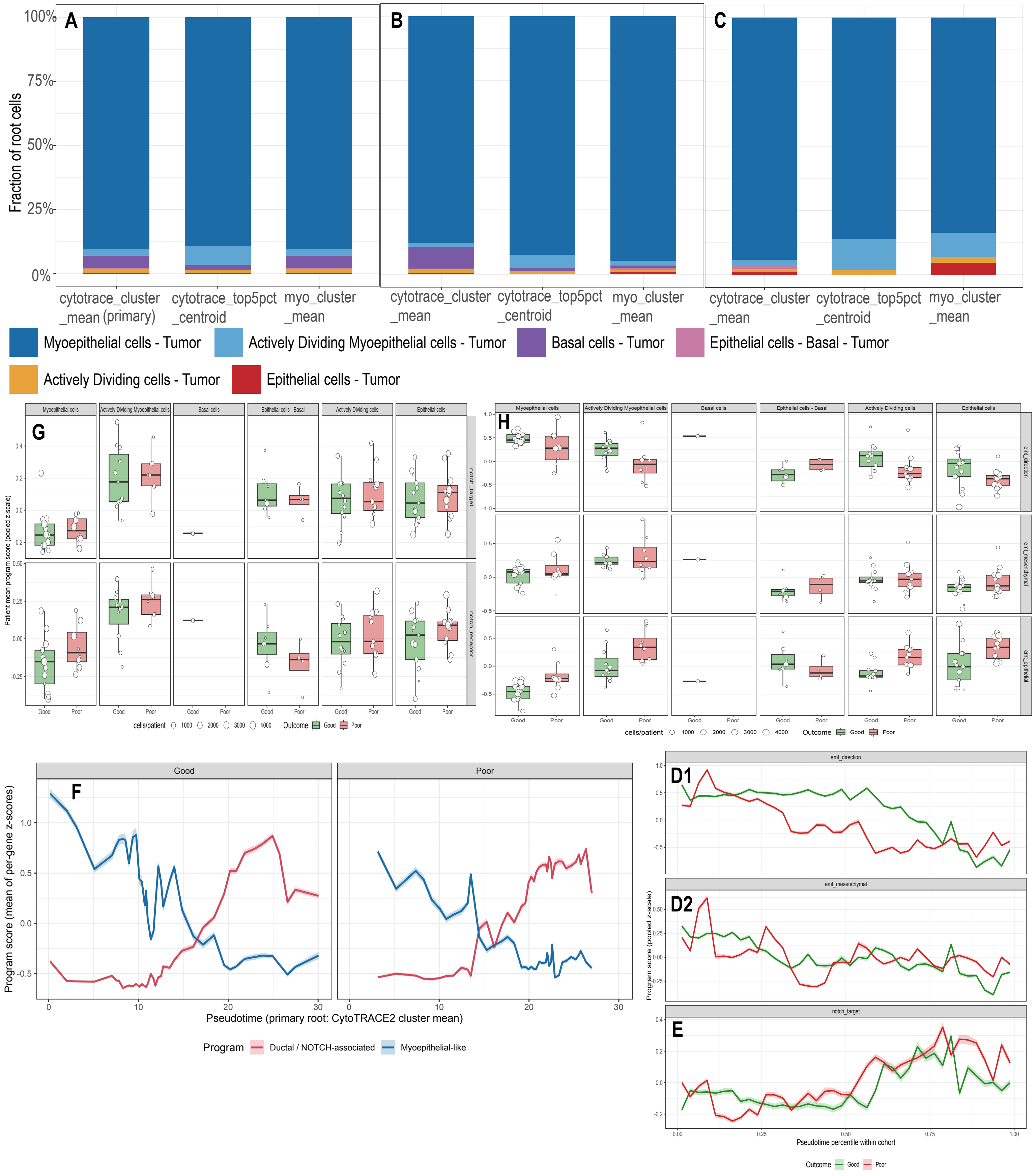

**Supplementary Figure Panel 4**

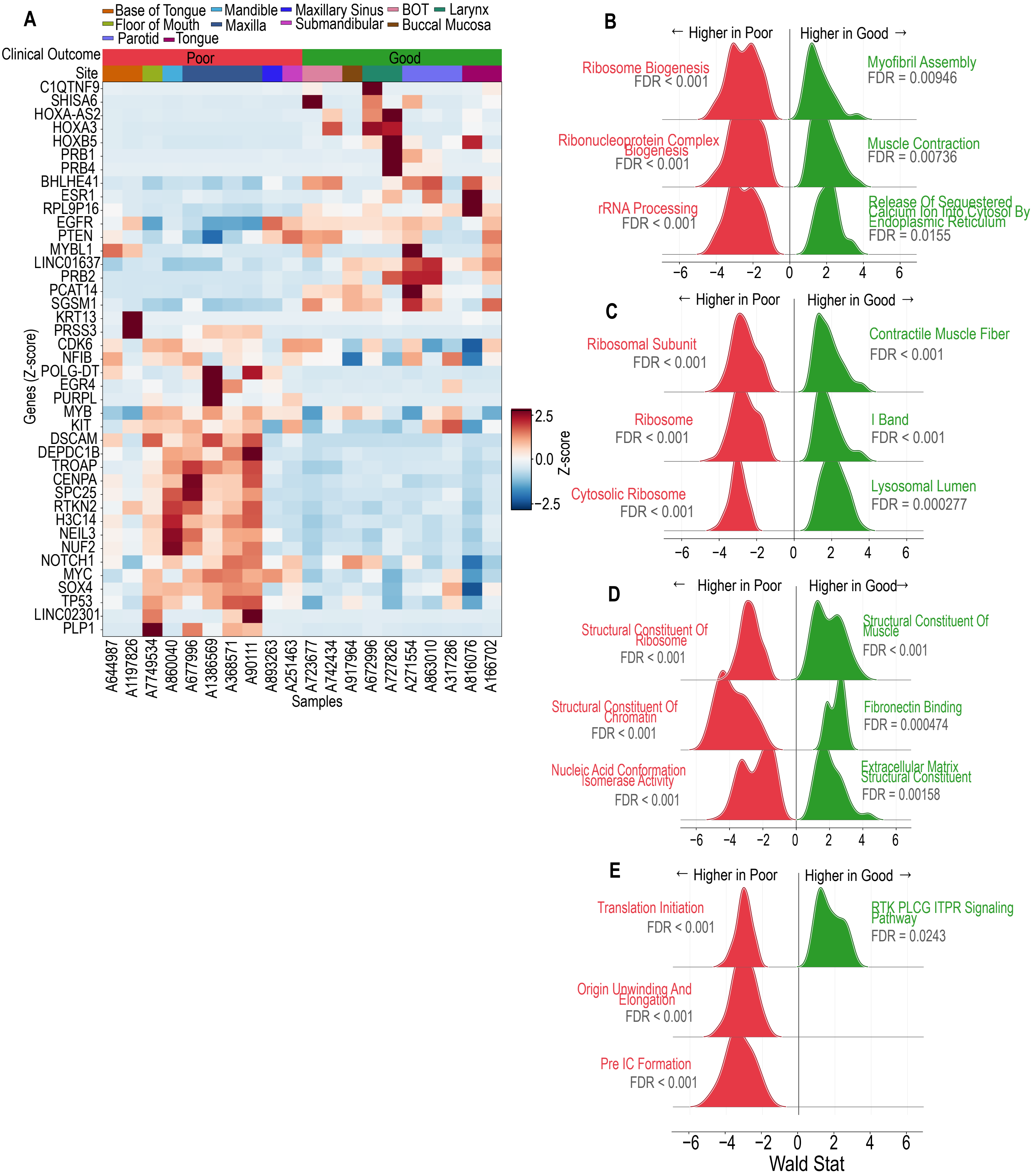

**Supplementary Figure Panel 5**

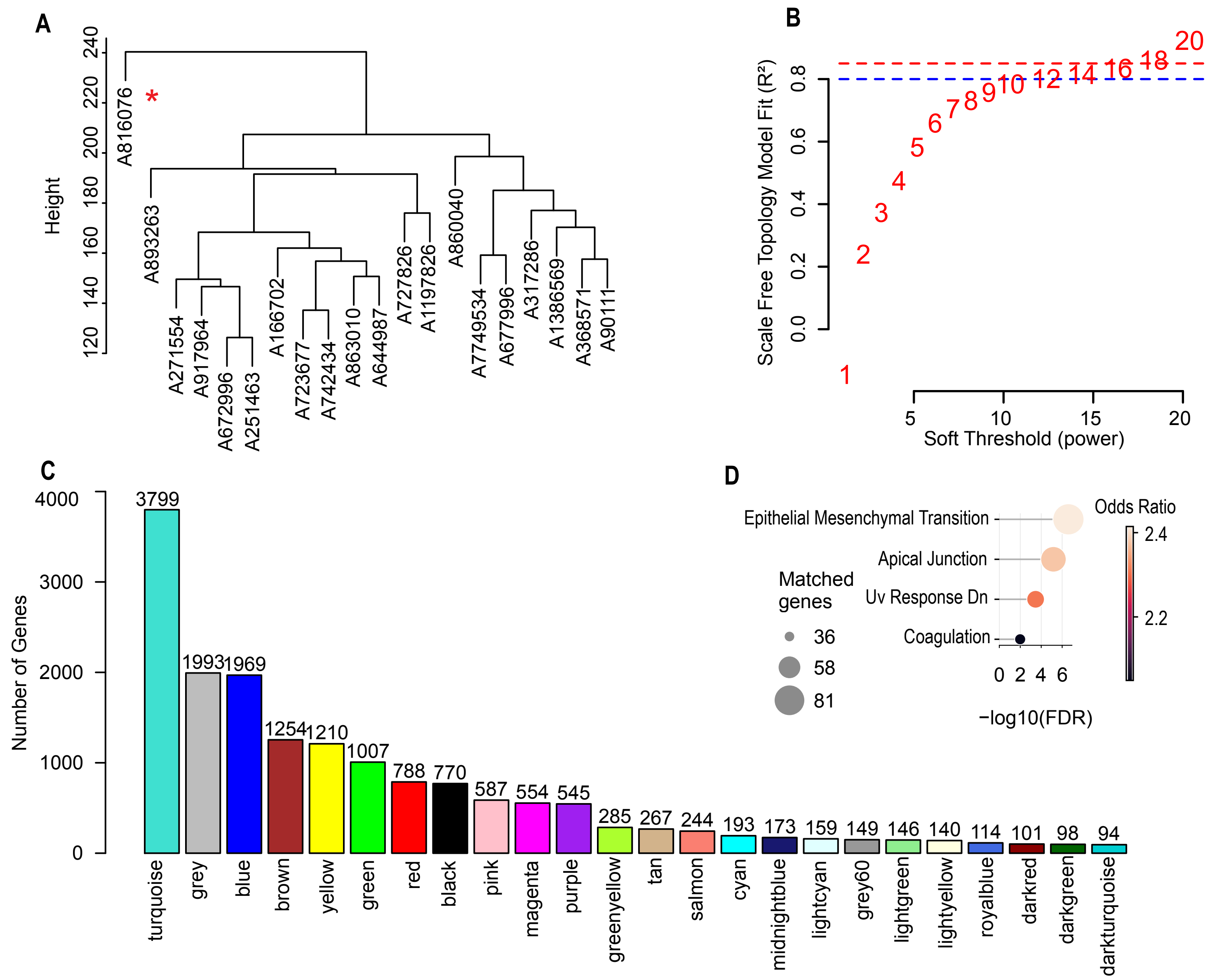

### Supplementary Figure Panel 6

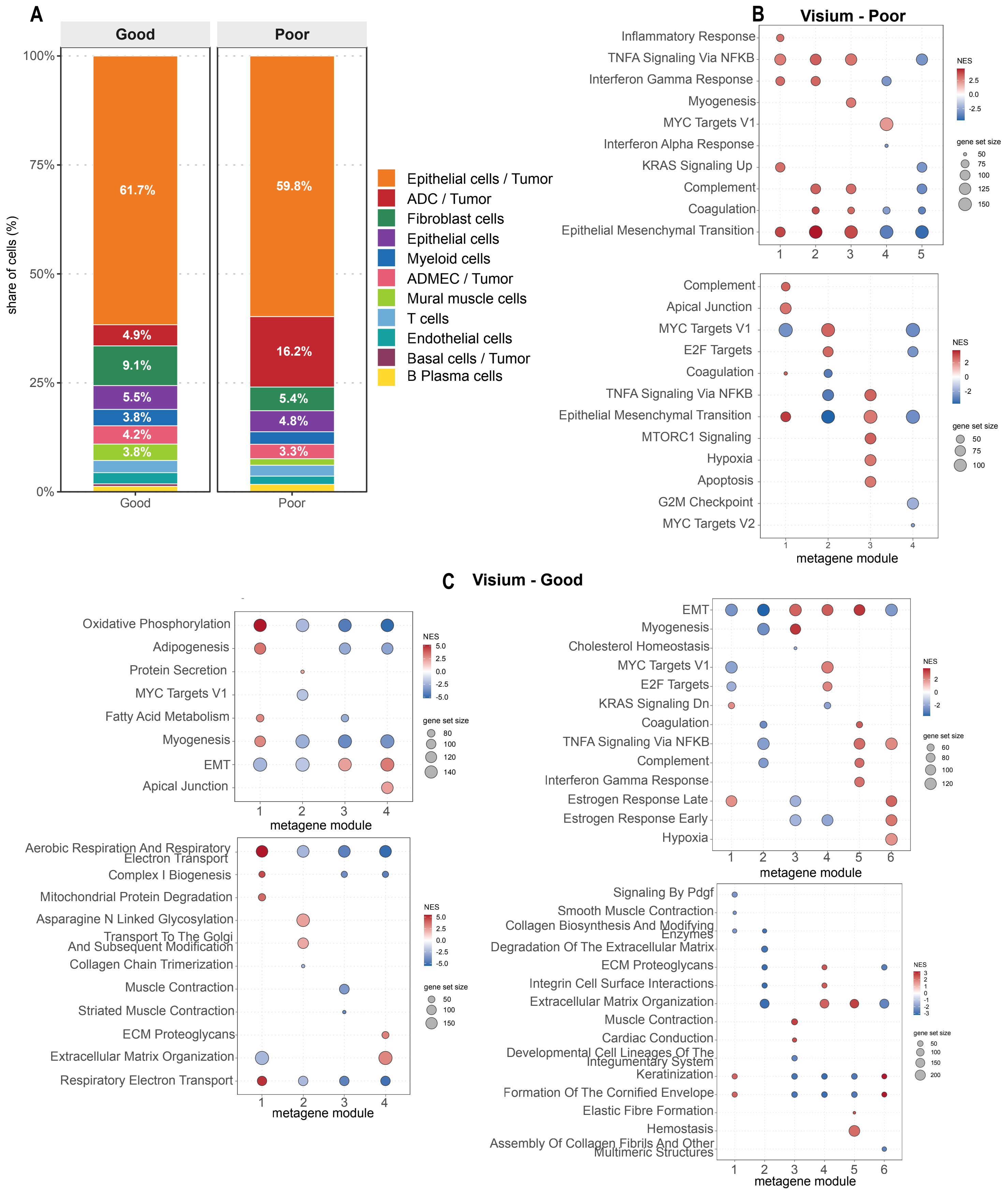

#### Supplementary Figure Panel 7

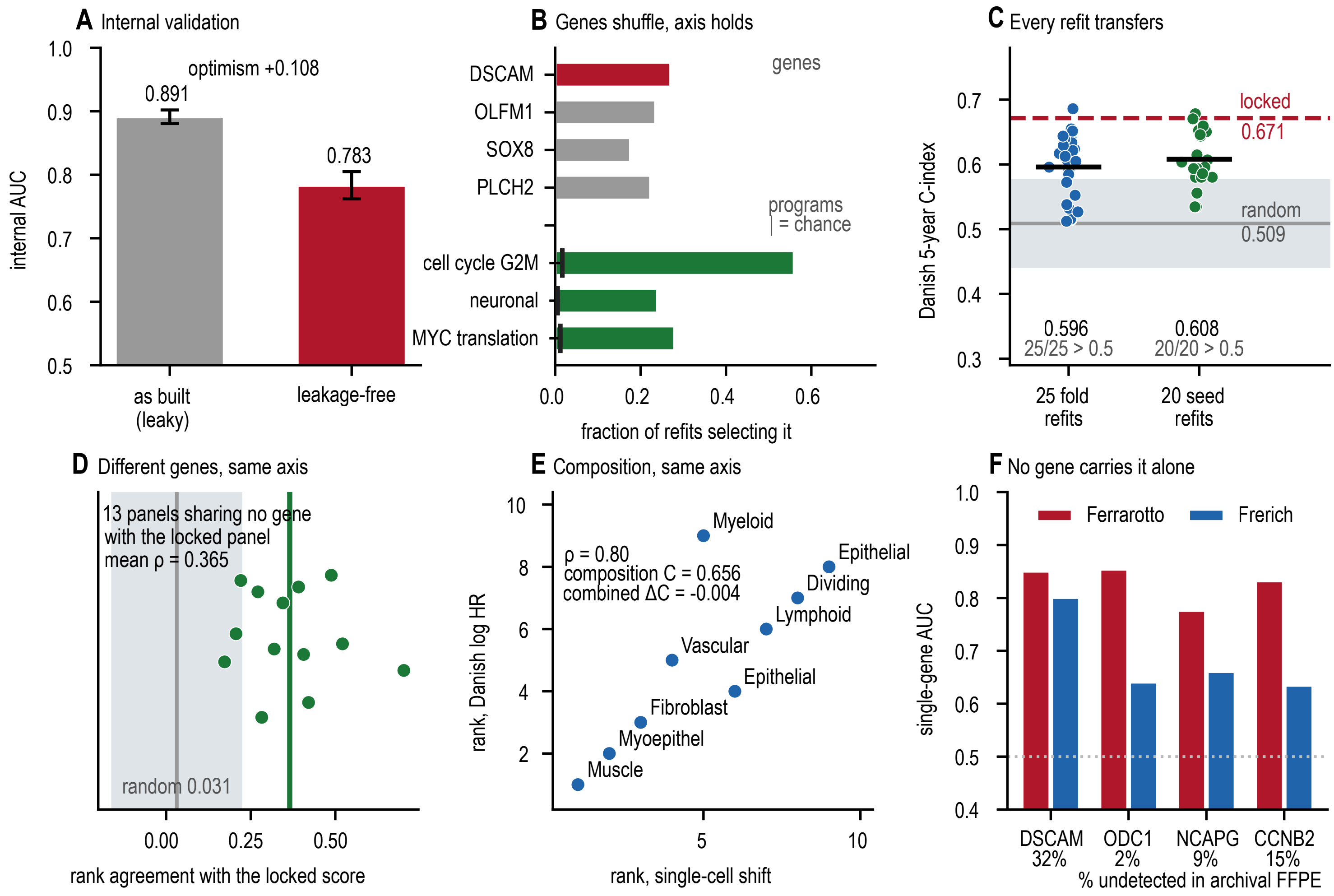

**Supplementary Figure Panel 8**

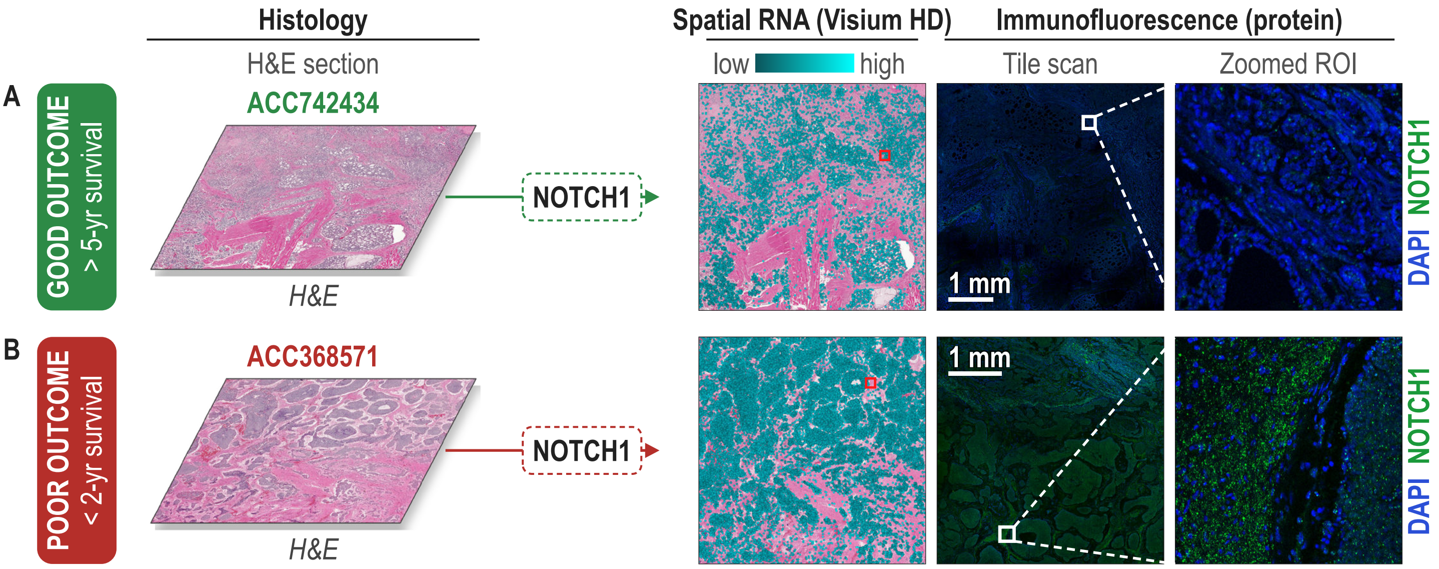

**Supplementary Information**

**Supplementary Table 1**

| Chemical | Supplier | Catalog number |
|---|---|---|
| semaphorin 3A | abcam | AB199475 |
| notch1 | abcam | AB52627 |
| ornithine decarboxylase ODC1 | abcam | AB193338 |
| c-myb | abcam | AB117635 |
| myc | proteintech | 60003-2-Ig |
| TrueBlack lipofuscin autofluorescence quencher 30x in DMSO | Biotium | 23011 |
| Vectashield Plus antifade mounting medium with DAPI | Vector Laboratories | H-2000-10 |
| Alexa Fluor 555 anti-mouse | Life Technologies | A21422 |
| Alexa Fluor 488 anti-rabbit | invitrogen | A11008 |

**Supplementary Table 2**

| | | | |
|---|---|---|---|
| TP53 | CDKN2A | CDKN2B | ATM |
| LATS1 | PLAGL1 | P73 | RB1 |
| CYLD | TSC1 | SUFU | KIT |
| C_KIT | EGFR | FGFR1 | FGFR2 |
| FGFR3 | FGFR4 | PDGFRA | PDGFRB |
| KDR | RET | PIK3CA | PTEN |
| AKT1 | MTOR | IGF1R | NOTCH2 |
| NOTCH3 | NOTCH4 | JAG1 | JAG2 |
| DLL1 | SPEN | FBXW7 | RBPJ |
| SMARCA2 | SMARCA4 | SMARCB1 | ARID1A |
| ARID1B | ARID4B | ARID5B | KDM6A |
| KDM5A | CREBBP | EP300 | KMT2C |
| KMT2D | BCOR | PBRM1 | BRD1 |
| BRD2 | EN1 | SOX4 | SOX11 |

| | | | |
|---|---|---|---|
| PRAME | SMO | CDK6 | KLF4 |
| FOXO1 | JUNB | FOSB | VGLL3 |
| ERBB3 | CTNNB1 | RUNX1 | FOXP2 |
| CDH2 | CDH5 | LIMA1 | CNTN6 |
| HSPG2 | SEMA3G | SEMA5A | FAT3 |
| FAT4 | BRCA1 | ATRX | UHRF1 |
| DCLRE1A | TXNIP | FGF16 | IL17RD |
| IDH1 | APC | NRAS | HRAS |
| BRAF | LRP1B | TERT | SF3B1 |
| MLH1 | MSH6 | MUC12 | TFAP4 |
| MAP10 | OTOP | RUNX3 | NBL1 |
| PRDM2 | NUP107 | MDM2 | |
